# Hypoxia induces a transcriptional early primitive streak signature in pluripotent cells enhancing spontaneous elongation and lineage representation in gastruloids

**DOI:** 10.1101/2021.07.21.452906

**Authors:** Natalia López-Anguita, Seher Ipek Gassaloglu, Maximilian Stötzel, Marina Typou, Iiris Virta, Sara Hetzel, René Buschow, Burak Koksal, Derya Atilla, Ronald Maitschke-Rajasekharan, Rui Chen, Alexandra L. Mattei, Ivan Bedzhov, David Meierhofer, Alexander Meissner, Jesse V. Veenvliet, Aydan Bulut-Karslioglu

**Affiliations:** Max Planck Institute for Molecular Genetics, Berlin, Germany; Charite Molecular Medicine, Berlin, Germany; Max Planck Institute of Molecular Cell Biology and Genetics, Dresden, Germany; Medical School of the Democritus University of Thrace, Greece; Bilkent University, Ankara, Turkey; Istanbul Technical University, Istanbul, Turkey; Max Planck Institute for Molecular Biomedicine, Münster, Germany; Department of Molecular and Cellular Biology, Harvard University, Cambridge, MA, USA; Department of Stem Cell and Regenerative Biology, Harvard University, Cambridge, MA, USA; Broad Institute of MIT and Harvard, Cambridge, MA, USA

## Abstract

The cellular microenvironment together with intrinsic regulators shapes stem cell identity and differentiation capacity. Mammalian early embryos are exposed to hypoxia in vivo and appear to benefit from hypoxic culture in vitro. Yet, components of the hypoxia response and how their interplay impacts stem cell transcriptional networks and lineage choices remain poorly understood. Here we investigated the molecular effects of acute and prolonged hypoxia on distinct embryonic and extraembryonic stem cell types as well as the functional impact on differentiation potential. We find a temporal and cell type-specific transcriptional response including an early primitive streak signature in hypoxic embryonic stem (ES) cells. Using a 3D gastruloid differentiation model, we show that hypoxia-induced T expression enables symmetry breaking and axial elongation in the absence of exogenous WNT activation. Importantly, hypoxia also modulates T levels in conventional gastruloids and enhances representation of endodermal and neural markers. Mechanistically, we identify Hif1α as a central factor that mediates the transcriptional response to hypoxia in balance with epigenetic and metabolic rewiring. Our findings directly link the microenvironment to stem cell function and provide a rationale supportive of applying physiological conditions in models of embryo development

## INTRODUCTION

In most early mammalian embryos, the first three cell types are established and maintained in oxygen tension ranging from 1.5 to 8% (Fischer and Bavister, 1993; Ottosen et al., 2006; Yedwab et al., 1976). These comprise the pluripotent epiblast, the primitive endoderm (PrE, also known as hypoblast), and the trophectoderm (TE). Upon differentiation, the extraembryonic PrE and TE give rise to the yolk sac and placenta, respectively (Rossant et al., 2003). Local oxygen levels in the embryo are likely subatmospheric until proper placentation at midgestation (Woods et al., 2018). Hypoxia is also endemic to adult stem cell niches (Mohyeldin et al., 2010) and solid tumors (Muz et al., 2015; Rankin and Giaccia, 2016) and as such is a common physiological component of the cellular microenvironment.

In vitro culture of pre-implantation mouse (Meuter et al., 2014; Nguyen et al., 2020), human (Dumoulin et al., 1999; Meintjes et al., 2009; Montfoort et al., 2020) and other mammalian embryos (Batt et al., 1991; Harvey et al., 2004) in hypoxia often improve embryo development or quality by e.g. increasing the inner cell mass size and reducing oxidative DNA damage (Houghton, 2021). Numerous studies on mouse and human ES cells revealed a beneficial effect on ES cell differentiation, especially towards endodermal lineages (Burr et al., 2017; Forristal et al., 2010; Houghton, 2021; Kusuma et al., 2014; Pimton et al., 2015). Hypoxia is also implemented in some protocols that model mammalian embryo development in a dish (Sozen et al., 2021), yet the mechanisms through which hypoxia exerts its beneficial effects are not clear. The severity and duration of hypoxia are major determinants of the hypoxic response, which is primarily executed by hypoxia-inducible factors (HIF). The canonical hypoxic response entails activation of glycolysis and angiogenesis genes by HIF1α (Muz et al., 2015; Podkalicka et al., 2020). In addition, hypoxia contributes to epithelial-mesenchymal transition (EMT) and invasiveness in various cancers (Muz et al., 2015; Rankin and Giaccia, 2016). EMT is also a cornerstone of embryonic development, as it enables cell movement and migration. In the mouse embryo, radial symmetry breaking happens at the time of gastrulation via formation of the primitive streak (Bardot and Hadjantonakis, 2020). The interplay between signaling pathways including WNT, BMP, and FGF and downstream activities of master transcription factors (TFs) such as Eomes, T (Brachyury), and Snail1 induce EMT and/or define mesodermal and endodermal cells on the posterior side of the embryo (Bardot and Hadjantonakis, 2020). Antagonization of these signals then defines the ectoderm on the anterior side (Bardot and Hadjantonakis, 2020).

Although WNT is not required for mouse preimplantation development (Fan et al., 2020), distinct WNT activities mediate long-term maintenance (Berge et al., 2011; Fan et al., 2020) and resolution of pluripotency (Bardot and Hadjantonakis, 2020) in vivo. In vitro, distinct pluripotent states can be captured (Kinoshita et al., 2021; Neagu et al., 2020; Ying et al., 2008) and differentiation programs can be mimicked (Brink et al., 2014; Hayashi et al., 2011) by modulating the activity of developmental signaling pathways including the WNT pathway. Exogenous WNT activation during differentiation of ES cells aggregates of a defined size leads to T expression and axial elongation, thus generating embryonic organoids resembling the post-implantation embryo (gastruloids) (Beccari et al., 2018; Berge et al., 2008; Brink et al., 2014; Turner et al., 2017; Vianello and Lutolf, 2020). Prolific WNT activity is however undesirable as it hinders the emergence of certain tissues including the neural lineages and reduces structural complexity (Girgin et al., 2021; Olmsted and Paluh, 2021). In vivo, transient, and fine-tuned gene activity underlies tissue diversification in the gastrulating embryo (Mittnenzweig et al., 2021). Moderation of pathway activity is thus crucial to achieve in vivo-like complexity in models of embryo development.

Here we investigated the molecular effects of acute and prolonged hypoxia on transcriptional networks and stem cell identities of embryonic and extraembryonic stem cells. We show that hypoxic ES cells are transcriptionally primed with a primitive streak signature including Wnt3 and T induction. This enables symmetry breaking in gastruloids without exogenous WNT activation. In conventional gastruloids, hypoxia enhances lineage representation in the presence of exogenous WNT activation. This transcriptional response is centrally regulated by Hif1α and also integrates metabolic and epigenetic responses.

## RESULTS

### Exposure of stem cells to hypoxia leads to cell-type-specific transcriptional responses

To investigate the impact of hypoxia on transcriptional programs of mouse early embryonic and extraembryonic stem cells, we used ES cells (Evans and Kaufman, 1981; Martin, 1981), extraembryonic endoderm stem (XEN) cells (Kunath et al., 2005) and trophoblast stem (TS) cells (Tanaka et al., 1998), which are in vitro models of the epiblast, PrE, and TE. The acute response to hypoxia in the form of HIF activation and metabolic adjustments occurs within minutes to hours (Pagé et al., 2002). However, the embryo might be exposed to hypoxia for about 9-10 days in vivo between fertilization and ∼E9-10 where proper placentation begins (Woods et al., 2018). Early embryonic and extraembryonic cells are thus likely exposed to hypoxia for at least a few days. To examine the effect of hypoxia on stem cell identity and function in a time-resolved manner, we cultured ES, TS, and XEN cells in hypoxia (2% O_2_) or normoxia (20% O_2_) for up to 7 days and first assessed stem cell marker expression, proliferation, and apoptosis rates (Figure 1A). In general, prolonged culture in hypoxia does not lead to increased cell death or morphological changes (Figure S1A, 1B). ES and TS cells proliferate slower in hypoxia (ES p=0.01, TS non-significant), with a slight increase of ES cells in the G_0_/G_1_ phase of the cell cycle (Figure S1B, S1C). XEN cell proliferation is largely not affected (Figure S1B). Importantly, ES cells remain undifferentiated and retain the pluripotency marker Nanog after 7 days of hypoxia exposure (Figure 1B, C). In contrast, hypoxia more drastically affects expression of stem cell markers in TS and XEN cells. Hypoxic TS cells downregulate Cdx2 expression and Gata4 expression is significantly increased in hypoxic XEN cells (Figure 1B, C). Thus, the response to hypoxia appears to be cell type-specific and may impact stem cell function in a tissue-specific manner.

**Figure 1.**
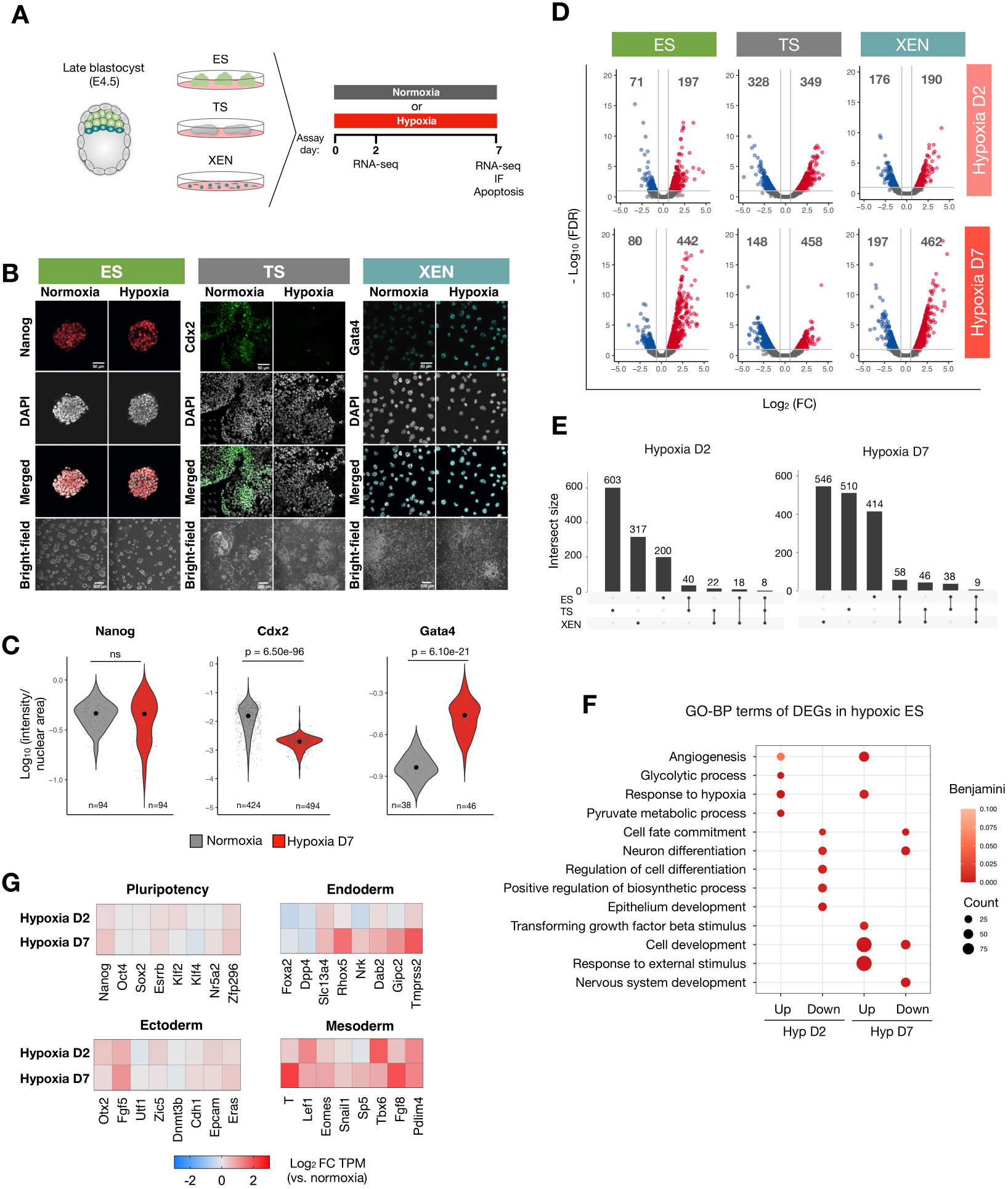
Hypoxia elicits cell type-specific transcriptional responses in ES, TS, and XEN cells. A) Experimental setup including a schematic of in vivo counterparts of the used stem cell types. B) Immunofluorescent (IF) staining of stem cell markers and bright-field images of ES, TS, and XEN cells cultured in normoxia or hypoxia for 7 days. Scale bars, 50 and 200 μm, respectively. C) Quantification of IF images including those displayed in (B). The fluorescent intensity of each nucleus was measured quantified and normalized to the nuclear area. Each dot represents a cell. N represents the number of cells quantified per sample. Statistical tests performed are unpaired two-sample Wilcoxon tests. D) Volcano plots depicting expression fold changes and significance on hypoxia day 2 and day 7 relative to normoxia (|FC|>1.5 and FDR >= 0.1 indicated by the gray lines). Red and blue indicate up- and down-regulated genes. Number of differentially expressed (DE) genes are indicated above each section. Two biological replicates were performed. See Table S1 for the full lists of DE genes. E) Overlap of DE genes across cell types on hypoxia day 2 and day 7. F) Gene ontology-Biological process (GO-BP) terms associated with DE genes in ES cells exposed to acute (d2) or prolonged (d7) hypoxia. Representative significant terms are shown (Benjamini-Hochberg adjusted p-value < 0.1). See Table S2 for full list of GO terms. GO terms associated with DE genes in TS and XEN cells can be found in Figure S1F-G and Table S2. G) Heatmaps showing expression levels of the indicated pluripotency and lineage-specific genes in hypoxic relative to normoxic ES cells. Germ layer markers were retrieved by analyzing RNAseq data of gastrulating mouse embryos published in (Grosswendt et al., 2020) as explained in Methods (see ‘Selection of germ layers and lineage markers’ section). TPM, transcripts per million. FC, fold change. See also Figure S1.

To probe the extent and specificity of the transcriptional response to hypoxia, we next profiled gene expression in each stem cell type on day 2 (acute) or day 7 (prolonged) of hypoxia exposure by bulk RNA sequencing (RNA-seq). Hierarchical clustering based on transcriptome profiles revealed primarily cell type-mediated clusters, indicating that hypoxia in general does not perturb cell identities (Figure S1D, S1E). In line with this observation, a moderate number of genes is significantly differentially expressed (DE) in each cell type and time point (|FC|>1.5, q-value <= 0.1, ES: 268 and 522 genes, TS: 677 and 606 genes, XEN: 366 and 659 genes on day 2 and day 7, respectively) (Figure 1D, Table S1). DE genes are largely cell-type-specific with minimal overlap (Figure 1E). Gene ontology analysis for biological processes of DE genes shows that the early response to hypoxia entails upregulation of glycolysis and angiogenesis genes in ES cells, but not TS and XEN cells (Figure 1F, Table S2), suggesting that the latter may be metabolically less dependent on oxygen (Figure S1F, S1G, Table S2). In all cell types, genes associated with cell development and differentiation are more deregulated on day 7, indicating temporally distinct effects of acute and prolonged hypoxia on transcription of specific gene subsets (Figure 1F, S1F, S1G). In ES cells, early mesoderm-instructive genes (E6.5-7.5) such as T, Tbx6, Fgf8, Tmprss2 are selectively upregulated in hypoxia (2-4 fold compared to normoxia), while ectoderm and pluripotency genes largely remain stable (Figure 1G). Genes that are later associated with node, notochord, and primitive gut development (Grosswendt et al., 2020) such as Krt19, Krt7, and Mixl1 are also upregulated, while those associated with brain and spinal cord development are downregulated. Thus, key mesoderm and endoderm lineage regulators appear to be specifically induced in hypoxic ES cells (Figure S1H). Taken together, stem cells present distinct temporal and cell type-specific transcriptional responses to hypoxia including lineage-specific genes. Importantly, these gene expression changes do not lead to spontaneous differentiation, they may however affect developmental trajectories later. Hereafter we focused on ES cells to investigate the hypoxia-associated activity of lineage-specific genes and its functional implications.

### WNT pathway genes are gradually upregulated during prolonged hypoxia

The induction of T, Eomes, Tbx6, Lef1 among other genes points towards increased WNT pathway activity in hypoxia (Figure 1G, S2A), which was previously observed in cancer cells and linked to epithelial-mesenchymal transition (EMT) (Muz et al., 2015; Rankin and Giaccia, 2016). Of all Wnt genes, Wnt3, Wnt4, Wnt7a, and Wnt7b are significantly upregulated in hypoxic ES cells (Figure S2B). Wnt3 activity at the posterior primitive streak mediates EMT during gastrulation by upregulating downstream targets such as T, Eomes, Tbx6, etc. (Bardot and Hadjantonakis, 2020). To investigate the temporal dynamics of the WNT pathway activity and its relation to oxygen availability, we next cultured ES cells in normoxia or varying degrees of hypoxia for up to 7 days and collected samples at regular intervals (Figure 2A-C). Wnt3 and T are gradually upregulated in hypoxia with an up to ∼4-fold increase over normoxia by day 7 (Figure 2B), while Eomes and Tbx6 are more modestly upregulated (∼1.5-fold). Pluripotency markers remain unchanged (Figure 2B, S2C). Importantly, Wnt3 and Eomes levels inversely correlate with oxygen availability, indicating association with an oxygen-mediated process (Figure 2C). Expression of pluripotency markers or oxidative phosphorylation genes does not correlate with oxygen levels, while glycolysis genes are expressed higher in more severe hypoxia (Figure S2D). Hypoxia-dependent induction of the Wnt pathway appears reversible, as expression levels of Wnt3, T, and Eomes revert to prior levels upon return of hypoxic cells to normoxia (Figure S2E). In hypoxia, WNT activity can be sustained for approximately 10 days, with Wnt3, T, Eomes, and Tbx6 transcript levels reverting to almost normoxic levels by day 16 (Figure S2E). HIF1α depletion in the nucleus on day 16 may underlie this pattern (Figure S2F).

**Figure 2.**
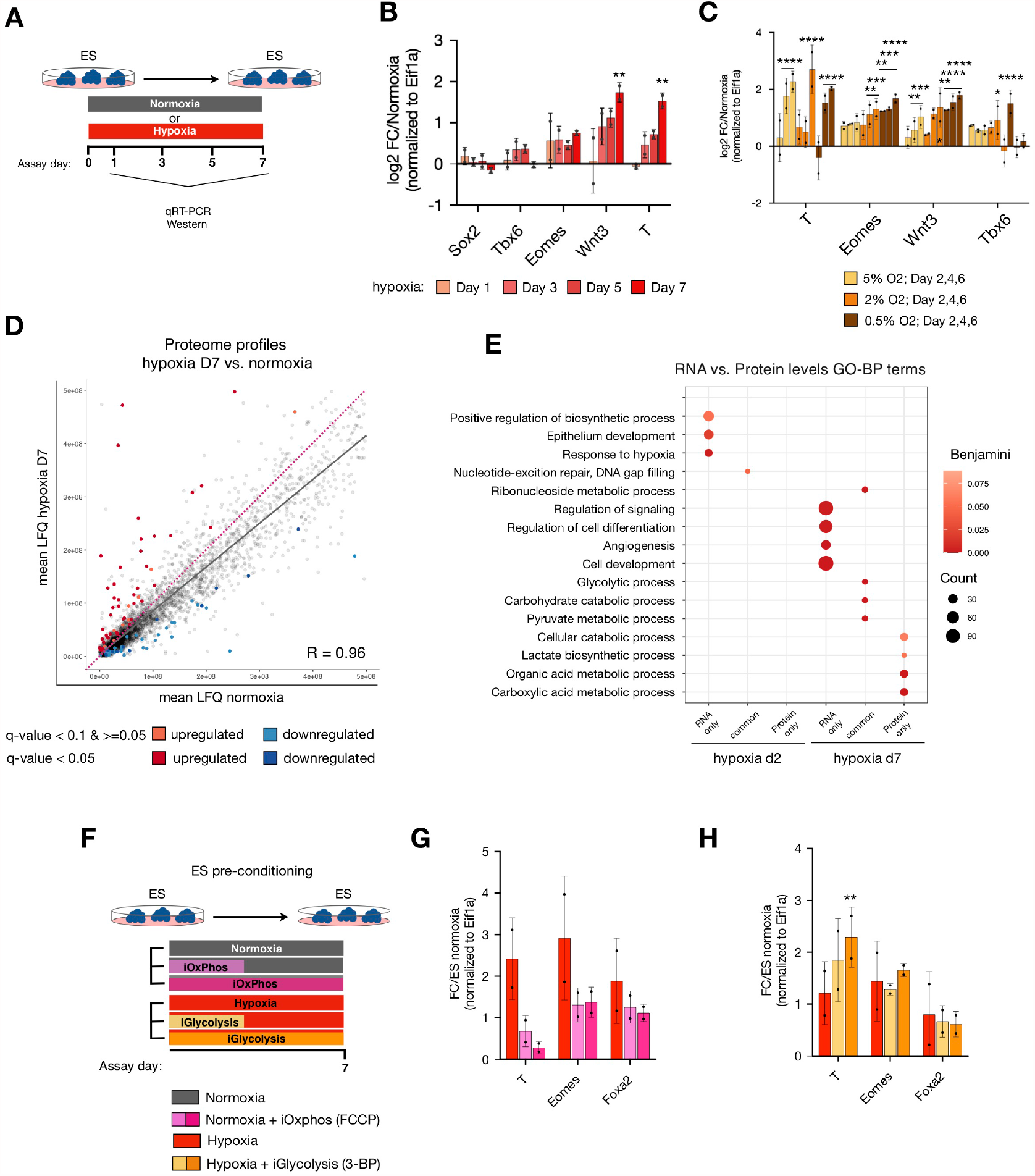
Transcriptional priming of ES cells towards differentiation in hypoxia by induction of WNT pathway genes. A) Schematic of experimental setup corresponding to panel (b). B) Relative expression levels of shown genes in hypoxic ES cells measured by RT-qPCR. Data represent log_2_FC over normoxia normalized to Eif1α and standard deviation for two biological replicates. Other tested genes are available in Figure S2C. The list of used primers is available in Table S10. The statistical test performed is a two-tailed paired Student’s t-test. C) Relative expression levels of shown genes in ES cells exposed to different levels of hypoxia measured by RT-qPCR. Data represent log_2_FC over normoxia normalized to Eif1α and standard deviation for two biological replicates. Other tested genes are available in Figure S2D. Statistical test performed is a two-way ANOVA. D) Proteome profiles of hypoxic (d7) vs normoxic ES cells. Colored dots indicate differentially expressed proteins (|FC|>1.5, q-value < 0.05 (dark red and blue) and q-value 0.05-0.1 (light red and blue)). Purple dotted line depicts the diagonal. LFQ, label-free quantification values. Complete proteome data and differentially expressed proteins (DEP) are available in Tables S3 and S4. E) GO terms associated with identified DE transcripts and proteins in hypoxia. Representative significant terms are shown (Benjamini-Hochberg adjusted p-value < 0.1). See Table S5 for full list of GO terms. F) Schematic of experimental setup. G, H) Relative expression levels of diagnostic genes in hypoxic ES cells vs normoxic ES cells treated with an oxidative phosphorylation inhibitor (G) or hypoxic ES cells treated with a glycolysis inhibitor (H). Data represent log_2_FC over DMSO-treated normoxic ES cells normalized to Eif1α and standard deviation for two biological replicates. See also Figure S2.

Oxygen depletion leads to curtailing of energy-dense processes including protein synthesis (Koumenis et al., 2002; Lee et al., 2020; Pettersen et al., 1986). To investigate whether differential expression at the transcript level is reflected at the protein level, we performed label-free mass spectrometry on ES cells in normoxia or exposed to acute or prolonged hypoxia (Figure 2D). Mass spectrometry identified 4260 proteins, of which 91.5% was also detected by RNA-seq (Figure S2G, proteome profiles are available in Table S3). In general, prolonged hypoxic ES cells retain a similar but globally downregulated proteome compared to normoxic ES cells (Spearman R=0.96, Figure 2D), with 63 and 43 proteins significantly up- or down-regulated (FC>1.5, q-value < 0.05, Table S4). Differentially expressed proteins (DEP) on day 7 of hypoxia are enriched for metabolic genes (Figure 2E, Table S4, S5). In contrast, development- and differentiation-associated genes are differentially expressed mostly at the transcript but not protein level (Figure 2E). This may be due to the limited sensitivity of mass spectrometry or alternatively due to selective translation or degradation of specific gene transcripts. Western blotting of whole-cell extracts showed low level but gradually increasing expression of T in hypoxic ES cells (Figure S2G), while Wnt3, Eomes, and Tbx6 could either not be detected or remained unaltered (data not shown). To assess whether T is functional under these conditions in ES cells, we analyzed the expression levels of T target genes (those bound and regulated by T during in vitro primitive streak differentiation (Lolas et al., 2014)) in hypoxic ES cells (Figure S2H). Several DE genes are found among putative T target genes but constitute a small fraction (4.25%). Furthermore, almost an equal number of putative T target genes are upregulated on day 2 and day 7. Since T induction at day 2 is modest (∼1.2-fold) at the transcript and protein level (Figure S2A, 2H), it is unlikely sufficient to directly activate downstream genes at that stage. In addition, putative T-activated genes are present among hypoxia-downregulated as well as hypoxia-upregulated genes (Figure S2H). We deduce that in ES cells hypoxia induces a transcriptional early primitive streak signature, which is not directly reflected at the protein level. As such, hypoxic ES cells do not resemble epiblast stem cells derived from the post-implantation embryos that comprise a mixture of T, FOXA2, and SOX2 expressing cells with anterior primitive streak characteristics (Kojima et al., 2014; Tsakiridis et al., 2014).

### Glycolysis counteracts the hypoxia-mediated early primitive streak signature

A major component of hypoxia response is the switch of cellular metabolism from oxidative phosphorylation to glycolysis. Indeed, metabolic rewiring constitutes the main hypoxia response in ES cells at the protein level (Figure 2E). Metabolic pathways not only determine ways of energy utilization but also impact cellular states (Hu et al., 2020; Rodriguez-Terrones et al., 2020). Therefore, we next set out to dissect the influence of glycolysis from other hypoxia-mediated events in the induction of early primitive streak genes. For this, we either induced glycolysis in normoxia (by inhibiting oxidative phosphorylation) or inhibited it in hypoxia acutely or continuously (Figure 2F). Induction of glycolysis in normoxia does not by itself induce T, Eomes, or Foxa2 expression and on the contrary decreases T expression (Figure 2G). Interestingly, inhibition of glycolysis in hypoxia further increases T expression (Figure 2H). Overall, T expression inversely correlates with the glycolytic state over time (Figure 2G, H). These results suggest that T levels in hypoxia are counteracted and balanced via a negative input of increased glycolysis, potentially enabling an early primitive streak-primed pluripotent state.

### HIF1α mediates the transcriptional priming in hypoxic ES cells

We next probed the primary hypoxia effector HIF1α for its potential regulation of early primitive streak genes. To test whether HIF1 activation suffices to upregulate these genes, we treated ES cells with IOX2 to inhibit the HIF1α destabilizer prolyl hydroxylase 2 (Figure 3A) (Chowdhury et al., 2013; Murray et al., 2010). IOX2 treatment thus stabilizes HIF1α in normoxia and allows to distinguish HIF1α-mediated and other effects of hypoxia. Treatment of ES cells with IOX2 resulted in the induction of Wnt3, T, and Eomes in a dose-dependent manner and at levels comparable to hypoxia (Figure 3A). Interestingly and similar to hypoxia, these genes are induced after prolonged treatment, while glycolysis genes are upregulated already on day 2 (Figure S3A).

**Figure 3.**
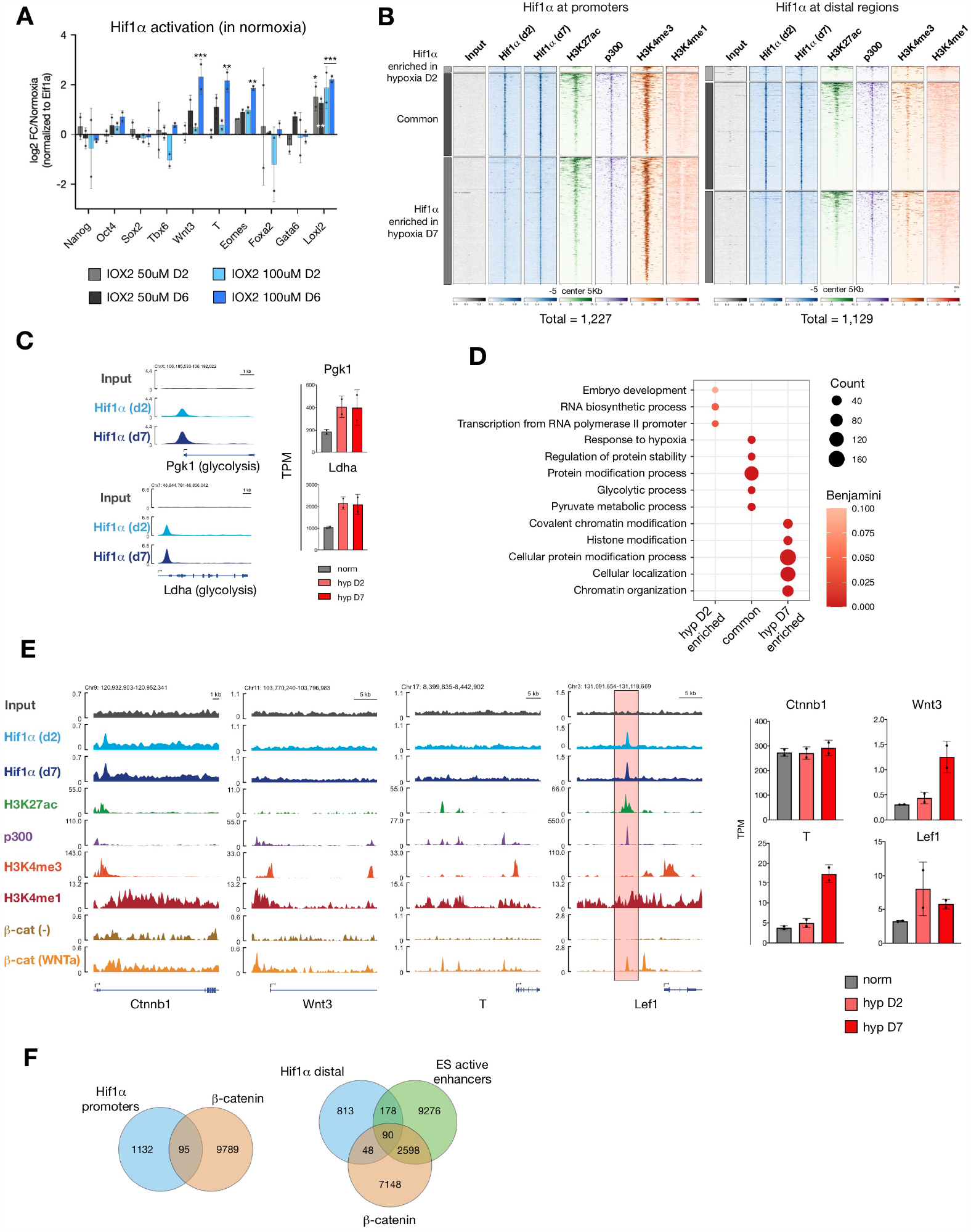
Hif1α mediates induction of WNT pathway genes in ES cells. A) Relative expression levels of shown genes in normoxic ES cells treated with the HIF1α activator IOX2 at indicated concentrations for 2 or 6 days. Data represent log_2_FC over DMSO-treated ES cells normalized to Eif1α and standard deviation for two biological replicates. Other tested genes are available in Figure S5A. The statistical test performed is two-way ANOVA. B) Density plots showing enrichments of the indicated genes and histone modifications at Hif1α-bound promoters and distal regions. ± 5 kb surrounding peak center is shown. Numbers below the density plots indicate the total number of HIF1α-bound sites at day 2 and day 7 of hypoxia. For a complete list of peaks, see Table S6. C) Genome browser views of HIF1α binding at the promoters of shown glycolysis genes. The right panel shows expression values of the shown genes in normoxic and hypoxic ES cells as measured by RNA-seq. D) GO-BP terms associated with HIF1α-target genes with HIF1α binding at promoters. Representative significant terms are shown (Benjamini-Hochberg adjusted p-value < 0.1). See Table S7 for the full list of GO terms. E) Genome browser views of the indicated genes and histone modifications at canonical WNT pathway genes (left). Expression values of the shown genes in normoxic and hypoxic ES cells as measured by RNA-seq (right). Red highlight depicts an active enhancer in ES cells. β-CATENIN binding data was retrieved from (Zhang et al., 2013). Histone modification and p300 binding data and the lust of ES active enhancers were retrieved from (Cruz-Molina et al., 2017). WNTa, WNT activation as employed in (Zhang et al., 2013). F) Venn diagrams showing the numbers of overlapping peaks with HIF1α and β-CATENIN potential co-binding at promoters (left) and ES active enhancers. See also Figure S3.

To investigate whether developmental genes are induced via direct HIF1α binding at promoters, we next profiled genomic binding sites of HIF1α by ChIP-seq on day 2 and day 7 of hypoxia treatment. HIF1α occupies promoters of 505 and 1,193 genes on day 2 and day 7, of which 471 are shared and include canonical targets such as glycolysis genes (Figure 3B-D, Table S6). In general, HIF1α continuously occupies glycolytic genes within the treatment time frame, while later-targeted genes are enriched for chromatin regulators (e.g., *Eed* and *Kdm4c*) and cellular localization genes (e.g., Prcc and Tpr) (Figure 3D, S3B, Table S7). However, HIF1α binding at non-metabolic genes does not directly affect transcriptional output (Figure S3B). Only 39 out of 1,193 genes with HIF1α enrichment at promoters on day 7 are significantly upregulated. We did not detect HIF1α binding at promoters of WNT pathway effectors except for a significant but minor enrichment at the β*-catenin* promoter, which does not result in altered expression (Figure 3E and S3C).

Notably, HIF1α also binds a comparable number of distal regions, some of which are enriched for H3K27 acetylation, H3K4me1, and p300 and are devoid of H3K4me3 in line with an enhancer signature (Figure 3B, Table S6) (Cruz-Molina et al., 2017). Although gene regulation by enhancers cannot be simply attributed to proximity in the linear genome, we find HIF1α binding at regions proximal to several genes including the WNT effector TF *Lef1* which is upregulated already on day 2 of hypoxia (Figure 3E, S3B). Lef1 is bound and activated by β-CATENIN upon WNT stimulation of ES cells but not in normal ES culture conditions (Figure 4E, compare β-catenin (WNTa) vs β-catenin (control), data from (Zhang et al., 2013)). HIF1α localization overlaps with the distal β-CATENIN binding site suggesting potential colocalization (Figure 3E). More generally, HIF1α and β-CATENIN share 95 target promoters and 90 target ES active enhancers (Figure 3F). We did not observe β-CATENIN stabilization or mainly nuclear localization in hypoxia as reported earlier (Mazumdar et al., 2010) (Figure S3D). These results suggest that HIF1α mediates the induction of early primitive streak genes in hypoxia but mostly not by direct DNA binding.

**Figure 4.**
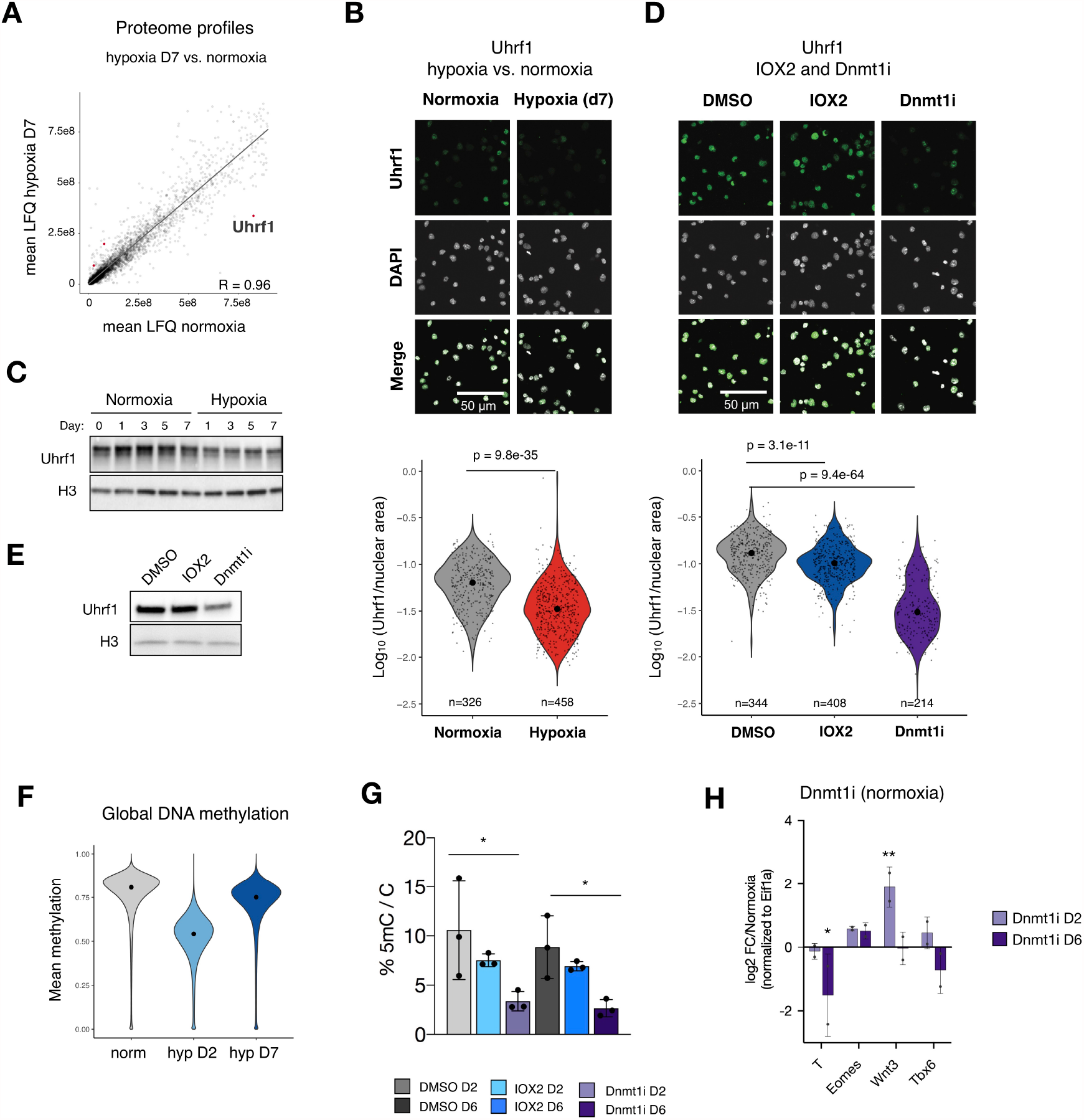
Depletion of Uhrf1 leads to global DNA demethylation in hypoxia. A) Proteome profiles of hypoxic (d7) vs normoxic ES cells. Epigenetic regulators that are significantly altered at the protein level are highlighted. LFQ, label-free quantification values. B) IF images and quantifications of Uhrf1 in hypoxic (d6) and normoxic ES cells. Fluorescent intensity of each nucleus was measured and normalized to nuclear area. Each dot represents a cell. N represents number of cells quantified per sample. Statistical tests performed are unpaired two-sample Wilcoxon tests. C) Western blots showing protein expression levels of Uhrf1 in normoxic and hypoxic ES cells. H3 was used as loading control. D) IF images and quantifications of Uhrf1 in normoxic ES cells treated with the HIF1α activator IOX2, a DNMT1 inhibitor, or DMSO as control. Fluorescent intensity of each nucleus was measured and normalized to nuclear area. Each dot represents a cell. N represents number of cells quantified per sample. Statistical tests performed are unpaired two-sample Wilcoxon tests. E) Western blots showing protein expression levels of Uhrf1 in normoxic ES cells treated with the HIF1α activator IOX2, a DNMT1 inhibitor, or DMSO as control (e). H3 was used as loading control. F) Global DNA methylation levels in normoxic and hypoxic ES cells determined by WGBS. Data represent mean of two biological replicates. G) DNA 5mC methylation levels in normoxic ES cells treated with the HIF1α activator IOX2, a Dnmt1 inhibitor, or DMSO as control measured by mass spectrometry. Data represent 5mC normalized to total cytosine (C) in three biological replicates. Statistical test performed is a one-way ANOVA. H) Relative expression levels of shown genes in ES cells treated with the a DNMT1 inhibitor for 2 or 6 days. Data represent log_2_FC over DMSO-treated ES cells normalized to Eif1α and standard deviation for two biological replicates. Statistical test performed is a two-way ANOVA. P value style: p>0.05 (ns), p<0.05 (*), p<0.01 (**), p<0.001 (***), p<0.0001 (****). See also Figure S4.

### Hypoxia leads to global DNA demethylation through depletion of UHRF1

Transcriptional control is exerted by TFs as well as modifiers of DNA and chromatin. While the hypoxic ES cell proteome is mostly stable, we noticed and confirmed the significant depletion of the DNA methyltransferase 1 (DNMT1) partner protein UHRF1 (Figure 4A-C). Its transcript levels remain unchanged, ruling out direct transcriptional control (Figure S4A). HIF1α activation in normoxia reduces UHRF1 levels, albeit to a lesser extent compared to hypoxia (Figure 4D, E), suggesting partial control by HIF1α. UHRF1 partners with DNMT1 for maintenance of DNA methylation during cell division (Bostick et al., 2007; Sharif et al., 2007). Although CG-rich gene promoters, including those of many developmental genes, are already depleted of DNA methylation, gene expression may be controlled by methylation of other regulatory elements such as enhancers (Feldmann et al., 2013; Stadler et al., 2011). Therefore, we investigated DNA methylation levels in ES cells exposed to acute (day 2) or prolonged hypoxia (day 7) by performing whole genome bisulfite sequencing (WGBS). Hypoxia exposure leads to a global decrease in DNA methylation, at a higher extent upon acute than prolonged exposure (32% vs 7%) (Figure 4F). We observe increased expression of the de novo DNA methyltransferase DNMT3B in the remethylation time frame (Figure S4B). Expression of the TET DNA demethylases does not change, although their activity might be altered due to the inherent oxygen sensitivity (Figure S4B, Table S3) (Burr et al., 2017). DNA methylation is comparably reduced at most genomic features as well as at differentially expressed genes (Figure S4C-E). HIF1α activation in normoxia via IOX2 treatment only moderately decreases DNA methylation, similar to the limited response of UHRF1 (Figure 4D, G). To probe whether DNA demethylation in normoxia can induce WNT pathway genes, we cultured normoxic ES cells with a DNMT1 inhibitor for 2 or 6 days (Figure 4H). DNA demethylation acutely induces Wnt3 but not downstream genes at the transcript level (Figure 4H). These results suggest a potential additive effect of HIF1α activity and DNA demethylation in hypoxia, which together with glycolysis shapes the transcriptional response at developmental genes.

### Hypoxic gastruloids spontaneously elongate in the absence of exogenous WNT activation

To functionally test whether the hypoxia-induced early primitive streak signature enables emergence of cell states and tissues that resemble and arise from the primitive streak in vivo, we employed the gastruloid model. Aggregation of ES cells upon withdrawal of self-renewal factors results in embryoid bodies that contain cells corresponding to the three germ layers. Transient treatment of embryoid bodies generated from defined, low numbers of ES cells with the exogenous WNT activator CHIR99021 (Chi) between 48h and 72h of culture induces symmetry breaking and elongation and self-organization of the body axis, with polarized T expression at the posterior end (Beccari et al., 2018; Berge et al., 2008; Brink et al., 2014). Since hypoxia induces a transcriptional early primitive streak signature including Wnt3 and T, we reasoned that it might enable spontaneous elongation in the absence of exogenous WNT activation (termed ‘-Chi’ hereafter). To test this possibility, we generated aggregates from 400 ES cells and scored T expression, size and elongation index (Figure 5A-D). While structures cultured in normoxia throughout the procedure do not express T or spontaneously elongate in our hands, pre-conditioning of ES cells in hypoxia prior to aggregation induced T expression and slight elongation (Figure 5B-D, S5A-B, compare HN to NN condition). Interestingly, exposure of structures to hypoxia only during differentiation led to stronger localized T expression and more pronounced spontaneous elongation in a subset of structures (Figure 5B-D, S5A-B, NH condition).

**Figure 5.**
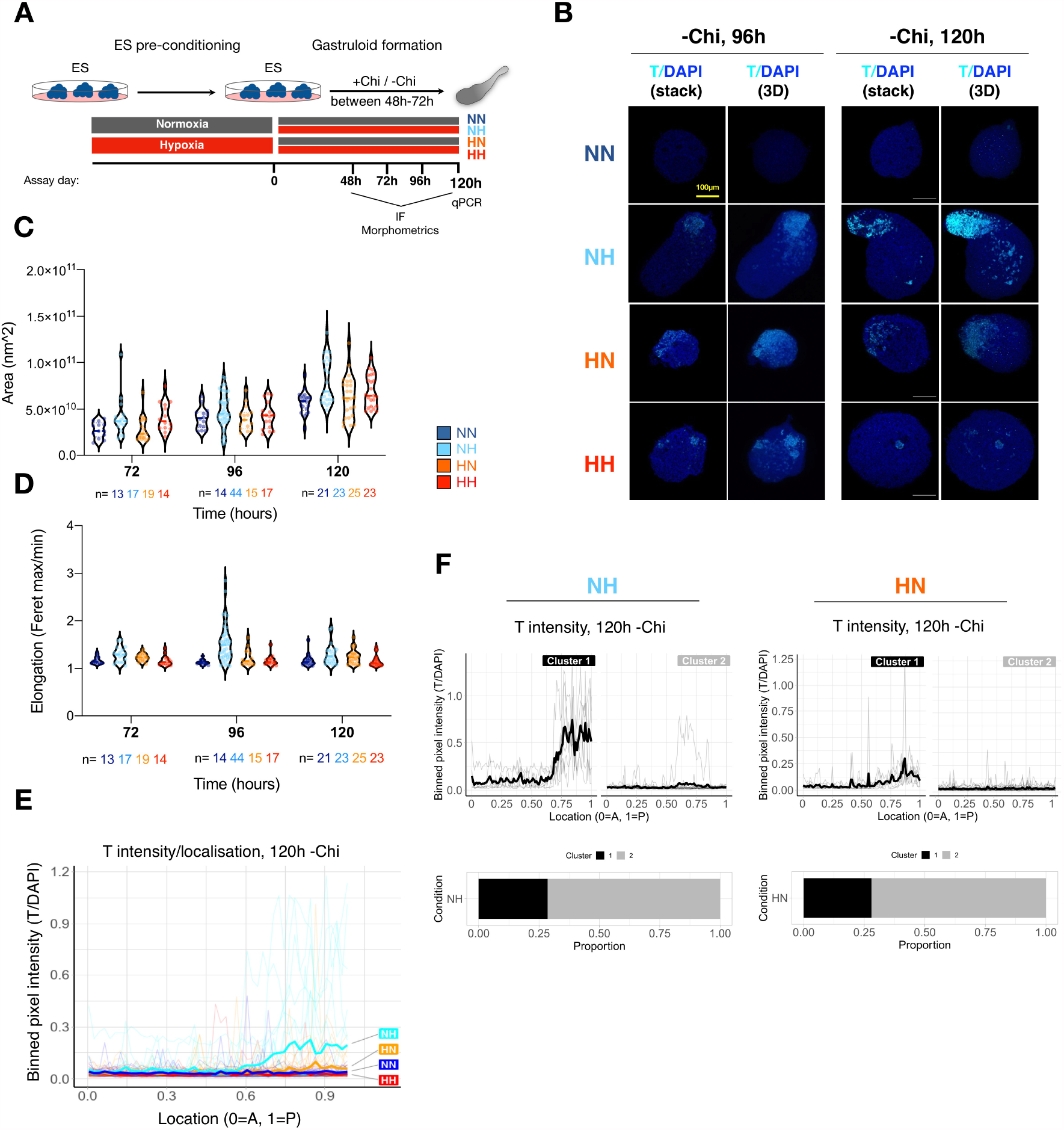
Hypoxia can induce spontaneous elongation of gastruloids in the absence of exogenous WNT activation. A) Schematic of the experimental setup. B) Confocal fluorescent microscopy images of representative -Chi gastruloids at 96h and 120h of culture. 3D, three-dimensional projection. C) Area of gastruloids at each time point and condition. Fluorescent images were used for quantifications. Each dot indicates a single structure and n indicates the number of analyzed structures at each time point and condition. D) Elongation index (defined as aspect ratio ferret max / feter min) of gastruloids at each time point and condition. Fluorescent images were used for quantifications. Each dot indicates a single structure and n indicates the number of analyzed structures at each time point and condition. Note the bimodality of the data, especially for the NH condition, with the structures with a higher index reflecting the proportion of gastruloids that spontaneously elongate. E) Localization of T signal along the anterior-posterior (A-P) axis of gastruloids in each condition at 120h of culture. T signal normalized to DAPI and binned at 1% length increments along each structure for plotting. See Methods for details. Thick lines show mean values and thin lines show data from individual structures. F) K-means clustering of the NH and HN structures presented in E) with n=2 clusters. See also Figure S5.

To further probe T expression, elongation dynamics, and variability in -Chi structures, we performed a time- and spatially-resolved quantitative image analysis of 284 structures obtained in four independent experiments (Figure 5E-F, S5C-D). We observe a dispersed T expression starting at 48 hours and posterior confinement starting at 72 hours after aggregation in NH and HN gastruloids (Figure S5B). This results in the spontaneous elongation of ∼30% of the NH and HN aggregates (vs 0% in NN condition, Figure 5F, S5D). T is most clearly induced and localized at the posterior end of the structure in the NH-Chi condition, although HN -Chi structures also present low level T expression (Figure 5E, F). These findings suggest that exposure to hypoxia equips ES aggregates with the capacity to self-initiate the developmental events characteristic of the post-implantation embryo, including symmetry breaking, polarization and axial elongation, pointing to the importance of the microenvironment in shaping cell commitment and tissue morphogenesis.

### Hypoxia modulates T expression in conventional gastruloids

Although -Chi HN and, in particular, NH gastruloids can spontaneously break symmetry, polarize and elongate, gastruloid formation efficiency and elongation capacity is limited under these conditions. To test the effect of hypoxia in a more robust model of embryo development, we next combined hypoxia with exogenous activation of WNT (termed ‘+Chi’ hereafter). In general, all +Chi structures elongate and show posterior T localization by 120h after aggregation (Figure 6A-B, S6A-C). Aggregation in hypoxia results in larger gastruloids (Figure 6B). Quantitative image analysis of 245 structures at 120h after aggregation across four independent experiments revealed that hypoxia moderates T expression, but does not alter localization (compare HN to NN, Figure 6C, D). This effect was more pronounced at 96h, when not only overall T expression was lower but also more localized under all hypoxic conditions (Figure S6D, E). Intriguingly, direct comparison of T localization and intensity in +Chi and -Chi gastruloids revealed that in the subset of -Chi NH gastruloids that spontaneously elongate, T levels and localization are similar to those observed in +Chi gastruloids at 120h (Figure 6E). In contrast, hyper-induction of T is seen in ∼15% of +Chi gastruloids but none of the -Chi NH gastruloids (Figure 4E). Notably, T expression dynamics resulting in its polarized expression are different in -Chi vs +Chi gastruloids. Whereas in -Chi gastruloids T is spontaneously induced in a small cluster of cells and becomes polarized at 72h; in +Chi gastruloids T is initially induced in most cells by 72h and is later polarized at 96h (compare Figure S5B to S6B). Taken together, the distinct mode of induction and expression dynamics of T might reflect the ability of hypoxia to provoke a physiological localized induction of T, whereas exogenous WNT induction might result in polarized T expression through both physiological and non-physiological phenomena.

**Figure 6.**
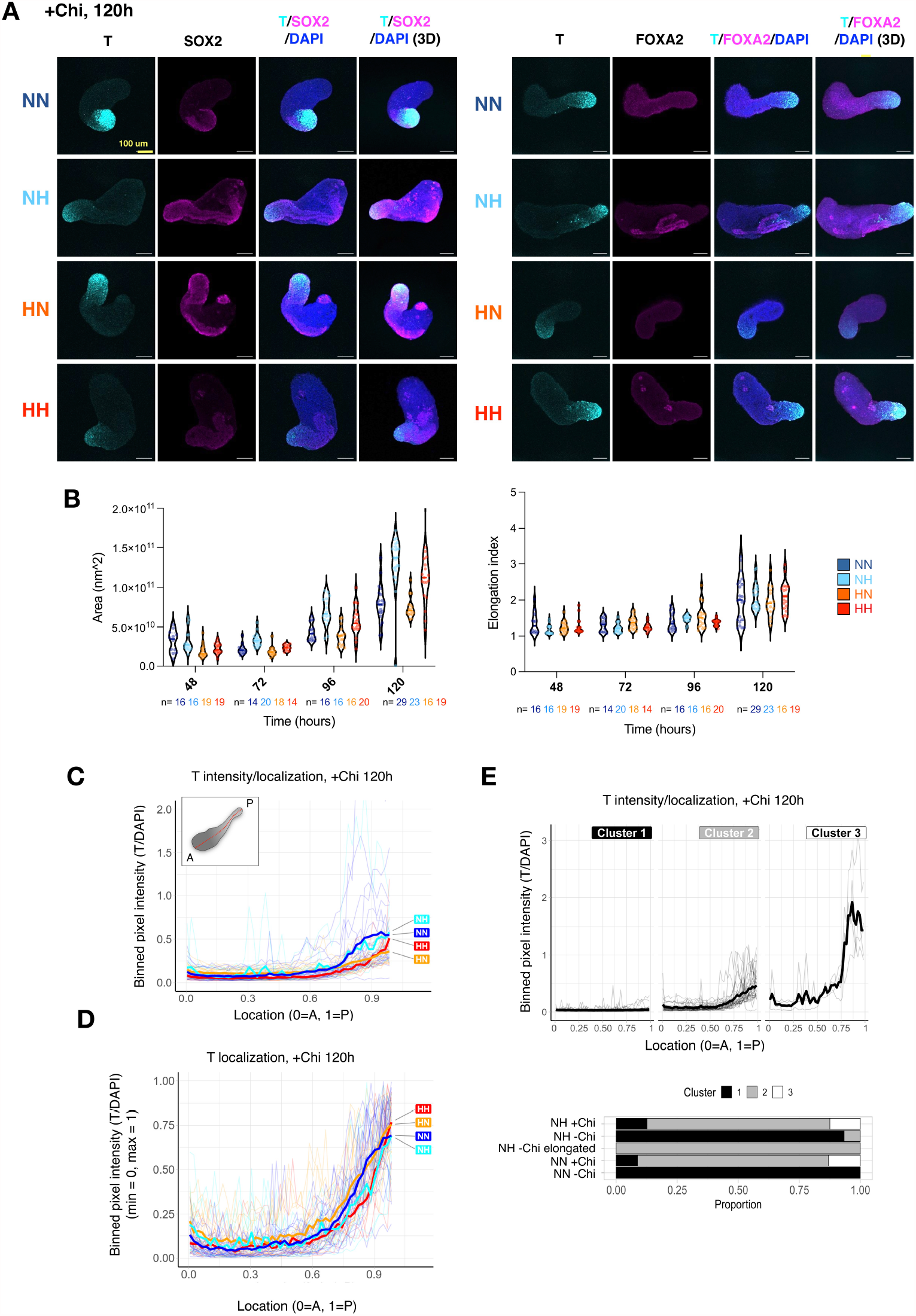
Hypoxia modulates T levels in conventional (+Chi) gastruloids. A) Confocal fluorescent microscopy images of representative +Chi gastruloids at 120h of culture. Images show a single z-stack, except those labeled 3D, which show 3D maximum intensity projections of the gastruloid. Additional structures are available in Figure S6C. B) Area and elongation index (defined as aspect ratio ferret max / ferret min) of +Chi gastruloids at each time point and condition. Fluorescent images were used for quantifications. Each dot indicates a single structure and n indicates the number of analyzed structures at each time point and condition. C, D) Localization of T signal along the anterior-posterior (A-P) axis of gastruloids in each condition. T signal was binned at 1% length increments along each structure for plotting. T intensity was normalized to DAPI stain, then was either plotted as such (C) or was further fitted in a 0-1 scale (D). See Methods for details. Thick lines show the mean and thin lines show data from individual structures. E) K-means clustering of all 120h +Chi structures plotted in C) with n=3 clusters. See also Figure S6.

### Enhanced representation of germ layer markers in conventional gastruloids cultured in hypoxia

Conventional +Chi gastruloids display localized T expression at the posterior end and at 120h contain a variety of cell states reminiscent of the post-occipital E8.5 embryo (Beccari et al., 2018; Berge et al., 2008; Brink et al., 2014; Turner et al., 2017; Vianello and Lutolf, 2020). However, the uniform application of a strong WNT signal in the conventional protocol is non-physiological, and might limit tissue diversification in gastruloids. To assess the relative abundance of cells representing the different germ layers in hypoxia, we stained +Chi gastruloids with SOX2 (as neuro-ectoderm marker), FOXA2 (as endoderm marker) in addition to T and analyzed their expression levels and patterns (Figure 6A, S6B-C, 7A-D). Conventional +Chi gastruloids cultured in normoxia throughout the procedure present substantial variability in expression patterns of SOX2 and FOXA2 (NN condition, Figure 6A, S6B-C). In our hands, SOX2 expression either covers the entire A-P axis or remains localized to the posterior end and FOXA2 expression remains minimal (Figure 6A, S6B, 7A-B). However, hypoxia exposure increases the proportion of SOX2^+^/T^-^cells as identified by single cell image analysis (Figure 7A). Remarkably, we identify FOXA2^+^/T^-^cells that self-organize around a lumen in a subset of the HN and NH gastruloids, reminiscent of development of the gut tube (Figure 7B, C). Quantitative expression analysis revealed that (pre)somitic mesoderm ((P)SM) marker genes Tbx6, Ripply2, Tcf15 and Uncx were also upregulated in gastruloids cultured under hypoxic conditions, independent on the ESC starting state (compare NH to NN and HH to HN in Figure 7D). In conclusion, these data show that i) hypoxic pre-conditioning of ES cells increases the induction of neuro-ectoderm in conventional gastruloids; ii) culture of gastruloids under hypoxic conditions enhance the levels of (P(SM)) and endodermal markers in conventional gastruloids; and iii) hypoxia can be sufficient to induce T expression and spontaneous elongation in -Chi gastruloids. Thus, hypoxia could potentially be harnessed in the culture of models of embryo development as a physiologically relevant means to induce axial elongation and improve germ layer representation.

**Figure 7.**
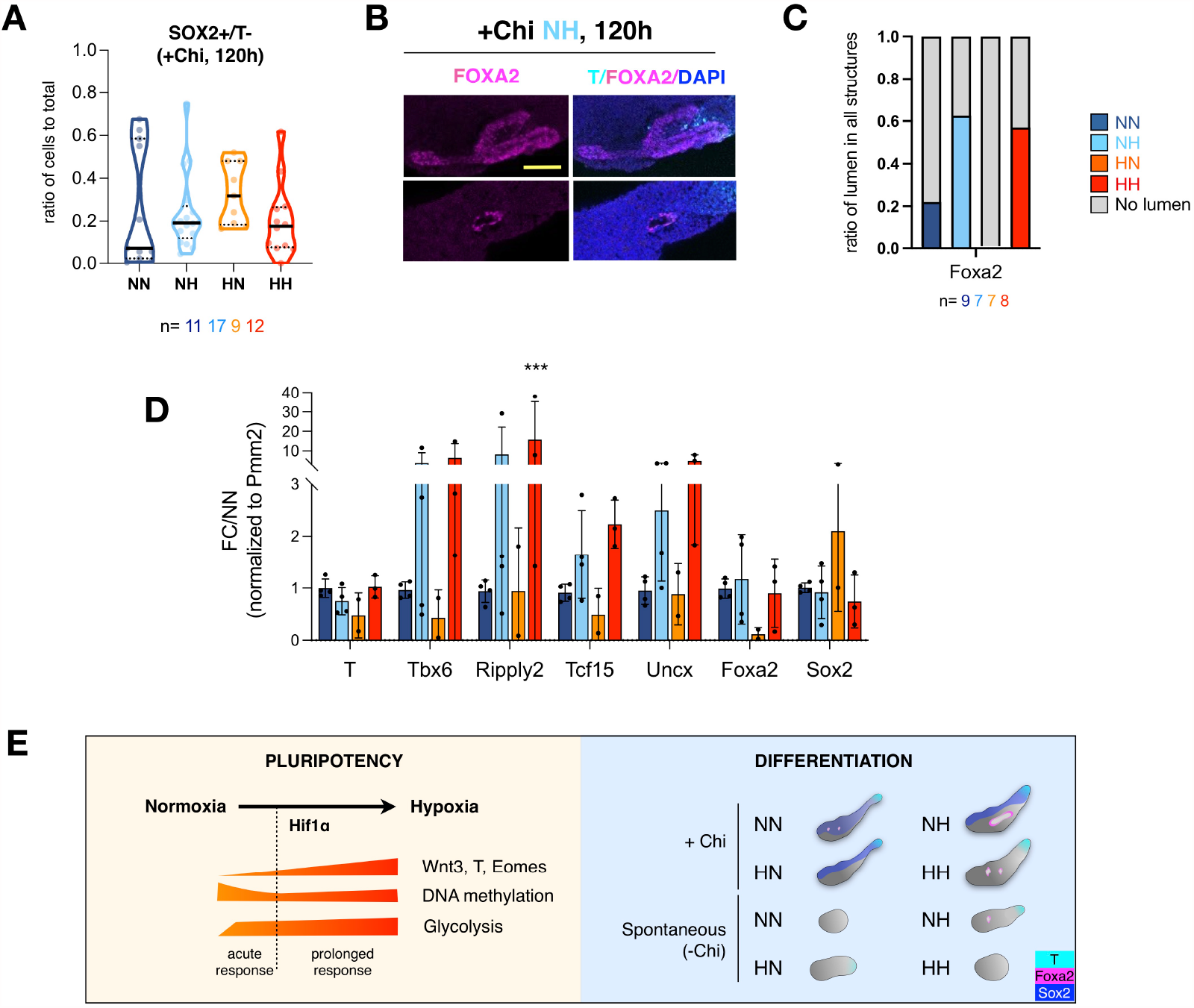
Hypoxia enhances lineage representation in conventional gastruloids. A) Proportion of SOX2 single positive cells in all quantified cells in each structure. Each gastruloid was quantified at the single-cell level for the presence of T/SOX2 staining. Each dot indicates a single structure. Thick lines indicate median and broken lines indicate quartiles. B) Confocal images of gastruloids showing gut tube-like structures. Scale bar is 100 μm. C) The ratio of structures containing a lumen surrounded by FOXA2 staining. n indicates number of analyzed structures in each condition. D) Relative expression levels of the indicated genes in normoxic or hypoxic gastruloids at 120h. Data represent FC over NN normalized to Pmm2 and standard deviation for 2-4 biological replicates. Each dot presents a biological replicate consisting of three pooled gastruloids. E) Working model of the components of acute and prolonged hypoxia response in ES cells and the outcome of exposure to hypoxia on differentiation of gastruloids.

## DISCUSSION

In this study, we present hypoxia as a critical microenvironmental factor shaping cell fate decisions in early stem cells. We show that hypoxia promotes a transcriptional early primitive streak signature in ES cells in a HIF1α-associated manner. In 3D gastruloids, hypoxia governs both cell fate decisions and tissue morphogenesis, leading to enhanced germ layer representation and spontaneous embryo-like morphogenetic events in the absence of exogenous WNT activation (Figure 7E). In extraembryonic stem cells, hypoxia induces transcriptional responses distinct from ES cells. Particularly, the significant decrease in Cdx2 expression in TS cells might favor a mural TE identity (Frias-Aldeguer et al., 2020). Interestingly, developmental genes appear to be responsive to hypoxia in all three cell types.

Whether this response also alters differentiation capacity of TS and XEN cells and how this would impact cellular crosstalk in co-culture systems are also of relevance to models of embryo development. We find that the ES cell epigenetic landscapes undergo major rewiring in response to hypoxia at least at the level of DNA demethylation. Additionally, several chromatin modifiers including Eed and Kdm4c show significant fluctuations at the protein level in response to hypoxia (Table S3, S4). Whether this results in further rewiring of the chromatin landscape and a potential interplay with DNA demethylation remains open. Increased glycolysis, a major component of hypoxia response, hampers the expression of lineage genes and counteracts Hif1-mediated induction of WNT pathway genes. In this context, glycolysis vs Wnt upregulation might achieve a balance in keeping transcriptionally-primed ES cells in the pluripotent state under hypoxic culture conditions.

Transcriptional induction of the WNT pathway in hypoxic ES cells is mediated by HIF1α activity, however, we do not detect direct HIF1α binding at promoters of *Wnt3, T, Eomes*, etc. Instead, HIF1α binds to active ES enhancers and could potentially co-localize with β-CATENIN. HIF1α binding at enhancers was previously observed in cancer cells (Andrysik et al., 2021; Platt et al., 2016). Here we show HIF1α binding at ES enhancers for the first time. Similarly, induction of various WNT responses by HIF1α has been widely documented in tumors and is associated with EMT and poor prognosis (Mori et al., 2016; Muz et al., 2015; Rankin and Giaccia, 2016; Schwab et al., 2012). Pluripotent and cancer cells are both fast-proliferating cells that share regulatory landscapes (Manzo, 2019). Therefore, it is conceivable that growth factors or regulators elicit similar responses in ES and cancer cells.

*Hif1*α KO is embryonic lethal at E9.5-10.5 due to neural tube defects and other embryonic abnormalities, together with placental defects largely attributable to defective angiogenesis (Kozak et al., 1997). Based on our data, these complex pathologies could indeed be rooted in events at much earlier stages of development. Mapping of lineage distribution in *Hif1*α KO embryos at gastrulation stages (E6.5-E9.5) at the single-cell level is required to reveal its control over lineage commitment. More broadly, HIF1α activity at canonical and noncanonical targets affects lineage choices in a cell type-dependent manner in health and disease (Allan et al., 2021; Lie et al., 2005; Shao et al., 2021). As such, hypoxia likely fine-tunes many developmental processes rather than altering developmental trajectories. Further understanding of the underlying mechanisms will inform strategies to harness this microenvironmental factor in models of embryo development.

The overall impact of hypoxia on development likely combines HIF1α-mediated and unrelated processes and encompasses altered epigenetic, transcriptional, and translational programs. For example, here we observe that most transcriptional changes are not reflected at the protein level, hinting at translational or post-transcriptional regulation. Translation has been shown to be globally suppressed in hypoxic cells (Koumenis et al., 2002; Pettersen et al., 1986). Translational profiling, e.g., via ribosome profiling, will likely identify selective translation as a regulator of ES self-renewal and differentiation under hypoxic conditions. On the other hand, part of the hypoxic response in ES cells is selective protein degradation as evidenced by gradual depletion of UHRF1 despite unaltered transcript levels (Figure 4A-C, Table S1). Which sequence features or protein associations mediate such selective degradation warrants further exploration. In addition to mediating DNA methylation, UHRF1 was shown to induce glycolysis, HIF1α expression, and EMT genes in cancers (Hu et al., 2019; Kim et al., 2017). We show here that UHRF1 is dynamically regulated in response to HIF1α activation and DNA demethylation (Figure 4D, E), however, whether UHRF1 levels influence glycolysis or EMT genes independent of DNA methylation remains to be further explored.

By testing the impact of hypoxia on lineage choices through gastruloid formation assays, here we show that hypoxia may act as a physiologically relevant means to induce symmetry breaking, polarization and axial elongation. Following these findings, hypoxia could be combined with a milder exogenous WNT activation to generate gastruloids that more closely mimic the embryo, and/or harnessed in other models of mouse post-implantation or human development (Moris et al., 2020; Olmsted and Paluh, 2021; Veenvliet et al., 2020; Xu et al., 2021). Particularly, the increased expression of (P)SM markers in hypoxic gastruloids suggests that hypoxia can be utilized to shift the decision making process of the bipotential neuromesodermal progenitors (NMPs) that contribute cells to both neural tube and somites in vivo (Koch et al., 2017; Tzouanacou et al., 2009). In trunk-like structures, in which neural tube and somite formation is unlocked by embedding gastruloids in low percentage matrigel, about 50% of the structures display a neural bias, with no clear somite formation, whereas in the other 50% formed somites are smaller than their in vivo counterparts and over time the propensity of NMPs to contribute to the somitic tissue vastly decreases (Veenvliet and Herrmann, 2021; Veenvliet et al., 2020). It will thus be of great interest to test if hypoxia can improve TLS formation.

The direct comparison of -Chi and +Chi gastruloids suggests that the developmental modes underlying T induction, symmetry breaking, polarization and axial elongation are distinct, with the non-uniform T induction in hypoxic -Chi gastruloids presumably more reliably recapitulating embryo dynamics. Interestingly, such non-uniform polarized T induction has been described in two recently published alternative gastruloid protocols, and in both cases leads to more complete models of embryo development, that include the cephalic tissues (Girgin et al., 2021; Xu et al., 2021). It will thus be interesting to see if cephalic tissues arise in hypoxic -Chi gastruloids.

Under our culture conditions, hypoxia has a clear endoderm-enhancing effect, in line with what has been reported for spontaneous and directed differentiation of mouse ES cells towards definitive endoderm (Pimton et al., 2015). The efficiency of endoderm induction in conventional gastruloids is currently subject to high variation. Whereas some studies have reported the formation of gut primordia in gastruloids, in other reports mainly neural and mesodermal tissue is formed (Beccari et al., 2018; Brink et al., 2020; Turner et al., 2017; Veenvliet et al., 2020; Vianello and Lutolf, 2020). Although the reason for this is unclear, the variation may be rooted in ES genetic backgrounds and/or pluripotency conditions, both linked to the propensity to form the different germ layers in gastruloids (Veenvliet and Herrmann, 2021). We therefore envision that hypoxia could be harnessed to more robustly induce gut primordia in gastruloids and other models of embryo development. Moreover, it will be of great interest to dissect how cell-intrinsic features (e.g. genetic background) interact with ESC pluripotency conditions and metabolic inputs (e.g. oxygen tension) in future studies.

Overall, our gastruloid data point to the importance of using physiologically relevant oxygen concentrations in models of embryo development. This is further corroborated by recent studies demonstrating that hypoxic culture conditions increase both the efficiency of the formation of blastocyst-like structures from human and mouse pluripotent stem cells, and ex utero culture of embryos from post-implantation to early organogenesis stages (Aguilera-Castrejon et al., 2021; Sozen et al., 2019, 2021). Thus, in addition to other modulators of the microenvironment such as medium composition, and matrix, we suggest oxygen tension to be taken into consideration when modelling developmental processes in a dish, or culturing mammalian embryos ex utero.

## ACKNOWLEDGMENTS

We thank members of the Bulut-Karslioglu Lab and Denes Hnisz for critical feedback, MPIMG sequencing, imaging and flow cytometry facilities for technical help and discussions. We thank Jennifer Shay for lab organization and assistance, Cordula Mancini for assistance, Beata Lukaszewska-McGreal for proteome sample preparation, Jalees Rehman (University of Illinois) for providing cell lines for initial characterization, Bernhard Herrmann for providing lab and office space, Thorsten Mielke and Beatrix Fauler for assistance with image acquisition. This project was supported by the Max Planck Society (D.M., A.M., J.V.V., A.B.-K.) a Bundesinstitut für Risikobewertung Bf3R grant No. 60-0102-01.P589 (J.V.V.) and the Sofja Kovalevskaja Award (Humboldt Foundation) to A.B.-K.

## AUTHOR CONTRIBUTIONS

A.B.-K. conceived the project. M.T. made initial observations and collected RNA-seq samples. S.I.G. performed the gastruloid experiments and analyzed the data together with R.B. and J.V.V. under the supervision of J.V.V.. M.S. performed IFs and TS and XEN characterization. I.V. performed metabolic analysis. A.L.M. prepared WGBS samples and S.H. performed WGBS analysis under the supervision of A.M.. B.K. and D.A. performed initial differentiation assays. R.C. helped with initial characterization of Hif1α pathway under the supervision of I.B. D.M. supervised mass spectrometry experiments. All other experiments and analyses were performed by N.L.A. with help from R.M.-R.. A.B.-K. supervised the project. J.V.V. and A.B.-K. wrote the manuscript with feedback from all authors.

## DATA AVAILABILITY

RNA-seq, ChIP-seq and WGBS data generated in this study has been deposited to the GEO database under the accession number GSE178628. Proteomics data has been deposited to PRIDE database under the accession number PXD026641. Full datasets will be released after publication.

## METHODS

### Cell lines and culture conditions

Wild-type E14 mouse ES cells (Sarah Kinkley Lab) were cultured without feeders on gelatin-coated plates (0.1% gelatin, Sigma-Aldrich G1393). The media was changed every day and cells were passaged every 2 days. At each passage, cells were dissociated using TrypLE (Thermo Fisher 12604-021) and replated at appropriate density. Cells were maintained at 37°C in a humidified 5% CO_2_ incubator. TS cells (Magdalena Zernicka-Goetz Lab) were cultured on feeders using TS cell culture media. Cells were passaged in a 1:10 to 1:20 ratio every 4 to 6 days. XEN cells (from Zernicka-Goetz Lab) were cultured on gelatin-coated plates using XEN cell culture media. Cells were passaged in 1:20 or 1:40 ratio every 2 to 3 days. Cells in normoxia were cultured with 20% O_2_ while cells in hypoxia were cultured with 2% O_2_ unless otherwise specified.

### Media and supplements

#### ES cell culture media

DMEM/High glucose with Glutamax media (Thermo, 31966047) supplemented with 15% FBS (Thermo, 2206648RP), 1x non-essential amino acids, 1x Penicillin/streptomycin, 1x β-mercaptoethanol, 1000U/mL LIF.

#### TS cell culture media

RPMI 1640 (Gibco, 61870010) supplemented with 1x Glutamax, 20% FBS, 1mM sodium pyruvate (Thermo, 11360070), 1.3x β-mercaptoethanol, 1x Penicillin/streptomycin, 25ng/ml FGF4 (R&D systems, 235-F4-025) and 1µg/ml Heparin (Sigma, H3149).

#### XEN cell culture media

DMEM + GlutaMAX (Gibco, 10566016) supplemented with 15% FBS, 1x non-essential amino acids, 1x Penicillin/streptomycin, 1x β-mercaptoethanol, 1mM Sodium pyruvate, 20mM HEPES (Sigma, H0887). Media was conditioned on mouse embryonic fibroblasts for 3 days and filtered before use. Then, feeder-conditioned media was mixed in a 7:3 ratio with unconditioned XEN media.

### Inhibitor treatments

For activation of Hif1α, cells were cultured in normoxia and treated with 50 or 100µM IOX2 (Sigma, SML0652) dissolved in DMSO (Sigma, D2650)

For DNA demethylation experiments, cells were cultured in normoxia and treated with 1µM Dnmt1 inhibitor (MedChem Tronica, HY-135146) dissolved in DMSO.

For inhibition of oxidative phosphorylation, cells were cultured in normoxia and treated with 1µM Carbonyl cyanide 4-(trifluoromethoxy)phenylhydrazone (FCCP) (Sigma, C2920) dissolved in DMSO.

For glycolysis inhibition, cells were cultured in hypoxia and treated with 50µM 3-bromopyruvate (3-BP) (Sigma, 376817) dissolved in DMSO.

### Growth curve

Cells were grown at a density of 0.25, 0.75, and 0.5 million cells per 10 cm dish for ES, TS, and XEN cells, respectively in normoxia or hypoxia 2% O_2,_ and were counted every day.

### Apoptosis assay

Cells were cultured in normoxia and hypoxia 2% O_2_ for 7 days. Then, the CellEvent Caspase-3/7 Green Flow Cytometry Assay Kit (Thermo, C10740) was used. Briefly, cells were collected and incubated with 500nM CellEvent Caspase-3/7 at RT for 1 hour. During the last 5 minutes, 1µM Sytox was added. As negative controls, cells were incubated in the absence of both reagents (double negative control), only with Caspase 3/7 or only with Sytox (single negative controls). Cells were analyzed on FACS AriaFusion. Data were analyzed using FlowJo (version 10) and plotted using GraphPadPrism (version 8).

### Cell cycle analysis

ES cells were cultured in normoxia and hypoxia 2% O_2_ for 7 days. To study cell cycle distribution, the Click-iT EdU Alexa Fluor 488 Flow Cytometry Assay Kit (Thermo, C10425) was used. Briefly, on day 7, cells were incubated at 37°C for 1 hour with 10µM EdU. Following the incubation, 1 million cells were collected and dislodged in fixative for at RT for 15 minutes. Then, cells were washed and mixed with the reaction cocktail (Alexa Fluor 488 azide) and incubated for 30 minutes. Cells were treated with 1:1000 DNA content stain FxCycle Violet in FACS media (PBS + 1% FBS). Cells were analyzed on FACS AriaFusion. Data were analyzed using FlowJo (version 10) and plotted using GraphPadPrism (version 8).

### RNA extraction and real-time qPCR

Total RNA was extracted using the Qiagen RNeasy kit (Qiagen, 74004), following the manufacturer’s instructions. 5 µg of RNA was used as input to generate complementary DNA (cDNA) with the High-Capacity cDNA Reverse Transcription Kit (Thermo, 4368814) using random primers. As a control, a reaction without reverse transcriptase enzyme (-RT control) was performed. cDNA was diluted 1:5 and was used to perform real-time quantitative PCR (qPCR) using primers designed to amplify specific target genes (Table S10) using KAPA SYBR FAST qPCR Master Mix (2X) ABI Prism (Thermo, KK4617) on the QuantStudio 7 Flex Real-Time PCR System (Applied Biosystems) thermal cycler. As a control, a reaction without cDNA was performed.

### Western Blot

#### Whole-cell extracts

Cell pellets were resuspended in RIPA buffer (Thermo, 89900) containing 1x protease inhibitor cocktail (Thermo, 78425), incubated at 4°C for 30 minutes, followed by centrifugation at maximum speed and 4°C for 20 minutes. Afterward, the supernatant was collected and the protein concentration was determined using Pierce™ BCA™ Protein-Assay (Thermo, 23225). 10-20 µg of protein was used for subsequent steps.

#### Subcellular fragmentation of cytoplasm and nucleus

Cell were washed with buffer A (10mM HEPES pH7.9 (Gibco, 15630-080), 5mM MgCl_2_ (Sigma, M8266), 0.25M Sucrose (Sigma, S7903), 0.1% Igepal 630 (Merck, 56741), 1x protease inhibitor cocktail, 1mM PMSF, 1mM NaVO_3_ and incubated on ice for 10 minutes. Afterward, samples were passed through 18G needles and centrifuged again for 10 minutes. Pellet corresponding to the nuclear fraction was resuspended in cold buffer B (10mM HEPES pH7.9, 1mM MgCl_2_, 0.1Mm EDTA (Jena Bioscience, BU-105), 25% glycerol (Sigma, G5516), 0.5M NaCl (Invitrogen, AM9760G) and 0.5% Triton X-100 were added and incubated on ice for 30 minutes. Samples were passed through 18G needles and sonicated using Bioruptor with the settings 30 seconds on 30 seconds off for 5 minutes. Subcellular fractions were quantified using Pierce™ BCA™ Protein-Assay. 10-20 µg protein was used for subsequent steps.

#### SDS-PAGE

Samples were mixed with 4x ROTI loading buffer (Carlroth, K929.2), heated at 98°C for 5 minutes, and loaded on 4–15% Mini-PROTEAN®TGX™ precast protein gels (Biorad, 4561083). Proteins were separated by electrophoresis at 70V for 15 minutes followed by 100V for approximately 1 hour using 10x Tris/Glycine/SDS running buffer (BioRad, 1610772).

#### Blotting and detection

Proteins were transferred to a PVDF membrane (Thermo, IB24001) using the iBlot 2 dry blotting system (Thermo, IB21001) and run at 20V for 7 minutes. After blotting, membranes were blocked with 5% milk in TBS-T buffer (Thermo, 28360) for 1 hour at room temperature. For detection of the protein of interest, membranes were incubated at 4°C overnight with primary antibody (Table S10) in 5% milk in TBS-T buffer, followed by secondary antibody at RT for 1 hour. For detection, membranes were incubated with ECL Western Blotting Substrate (Thermo, 32106) for 1 minute prior to imaging with the ChemiDoc system (BioRad).

### Immunofluorescence (IF)

IF was either performed on colonies or seeded single cells. For imaging of colonies, cells were cultured on Falcon 8-well Culture Slides (Corning, 354108) coated with gelatin (XEN, ES) or uncoated (TS). For single-cell staining, 50.000 to 100.000 cells were seeded on Matrigel (Corning, 356230) for 30 minutes. Cells were washed with 1xPBS before fixation using 4% PFA (Sigma, P6148) for 10 min at RT. After fixation cells were washed with 1xPBS and permeabilized using 0.4% Triton-X100 in PBS-T for 15 min at RT. After permeabilization, samples were washed in PBS-T and blocked in PBS-T + 2% BSA (NEB, B9000) + 5% donkey/goat serum (Jackson Immunoresearch/Dianova, 017-000-121) for 1h at RT. Cells were incubated with primary antibodies diluted 1:200 to 1:1000 in blocking buffer overnight at 4°C. Samples were washed three times 10 min at RT in PBS-T + 2% BSA and subsequently incubated with secondary antibodies diluted 1:1000 in blocking buffer for 1h at RT. Samples were washed three times for 10 min at RT in PBS-T + 2% BSA. Cells were mounted in Vectashield (H1200) with DAPI (Biozol, VEC-H-2000-10). Imaging was performed with confocal laser scanning microscope LSM800 (Zeiss). Fluorescence intensities were quantified using CellProfiler (https://cellprofiler.org/) (Carpenter et al., 2006). Intensities were normalized to the nuclear area and plotted in R using ggplot2 (Wickham, 2016).

### RNA-sequencing

#### Sample collection

Total RNA was extracted using the Qiagen RNeasy kit and quantified using Qubit 3.0. Two biological replicates were collected per condition.

#### Library preparation and sequencing

Library preparation was performed using KAPA RNA HyperPrep Kit (Kapa Biosystems, KR1350) (input amount 500ng RNA, adapter concentration 1.5 µM and PCR cycle number = 10)), and samples were sequenced using Illumina HiSeq4000, in 75-bp, paired-end format.

#### Mapping and analysis

Raw reads were subjected to adapter and quality trimming with cutadapt (version 2.4; parameters: --nextseq-trim 20 --overlap 5 --minimum-length 25 -- adapter A G A T C G G A A G A G C - A AGATCGGAAGAGC), followed by poly-A trimming with cutadapt (parameters: --overlap 20 --minimum-length 25 --adapter “A[100]” -- adapter “T[100]”). Reads were aligned to the mouse reference genome (mm10) using STAR (version 2.7.5a; parameters: --runMode alignReads --chimSegmentMin 20 -- outSAMstrandField intronMotif --quantMode GeneCounts) and transcripts were assembled using StringTie (version 2.0.6; parameters: -e) with GENCODE annotation (VM19). Table of read counts was generated using featureCounts function of the Rsubread package (version 1.32.4). To filter lowly expressed genes, count-per-million (CPM) was calculated using EdgeR package (version 3.26.8). Genes having > 0.4 CPM in at least 2 out of 6 samples were kept. Additionally, only coding protein genes were used for further analysis. Transcripts Per Kilobase Million (TPM) was used to generate normalized expression values. Differentially expressed genes (DEG) were determined using EdgeR package, applying multiple-testing adjusted P (FDR) <= 0.1 significance threshold and absolute fold-change > 1.5. Complete lists of DE genes are available in Table S1.

Principal component analysis was performed with the built-in R functions prcomp and hierarchical clustering was made with dendextend library (version 1.14.0) using dist function to compute distance between sample and hclust to perform hierarchical clustering. PCA and volcano plots were generated using ggplot2 (version 3.3.0). Heatmaps were generated using GraphPadPrism (version 8).

#### Selection of germ layers and lineage markers

Marker genes from different cell states were selected following the single-cell transcriptional reference profile of early post-implantation development from (Grosswendt et al., 2020). Early ecto-, endo- and mesoderm germ layers represents marker genes that were reliably assigned to cell states within these three lineage cell fates from E6.5 to E7.5.

For later cell states, lineage specific marker genes from node & notochord (E7.0-8.0, mesoderm lineage), neuromesodermal (E7.5-8.0, mesoderm lineage), paraxial & posterior mesoderm (E7.5-8.5, mesoderm lineage), primitive gut (E7.5-8.5, endoderm lineage) and for mid brain & spinal cord (E7.5-8.5, ectoderm lineage) were selected.

#### Gene ontology-biological process (GO-BP) analysis

GO-BP analyses were performed using DAVID web-tools (Dennis et al., 2003). Selected terms were plotted in dot plot format using ggplot2. Complete lists of GO terms are available in Table S2.

#### Analysis of T target genes

T ChIP-seq and RNA-seq data were retrieved from (Lolas et al., 2014). Briefly, T-activated (FC>2) or T-repressed genes (FC<-2) in *in vitro* primitive streak differentiated cells on day 4 vs. ES cells were selected. Expression levels of these T target genes in hypoxia day 2, day 7 and normoxia was plotted in heatmap format using pheatmap package (version 1.0.12) showing row z-score normalized expression values. Annotation bars were added to show which of these genes are differentially expressed in hypoxia day 2 and/or day 7 compared to normoxia.

### Global proteomics

#### Sample preparation

Proteomics sample preparation was done according to a published protocol with minor modifications (Kulak et al., 2014). In brief, 5 million cells in biological duplicates of 2 days and 7 days hypoxia treatment and normoxia controls were lysed under denaturing conditions in 500 µl of a buffer containing 3 M guanidinium chloride (G d m C l), 1 0 m M t r i s (2 - carboxyethyl)phosphine, 40 mM chloroacetamide, and 100 mM Tris-HCl pH 8.5. Lysates were denatured at 95°C for 10 min shaking at 1000 rpm in a thermal shaker and sonicated in a water bath for 10 min. 100 µl lysate was diluted with a dilution buffer containing 10% acetonitrile and 25 mM Tris-HCl, pH 8.0, to reach a 1 M GdmCl concentration. Then, proteins were digested with LysC (Roche, Basel, Switzerland; enzyme to protein ratio 1:50, MS-grade) shaking at 700 rpm at 37°C for 2 hours. The digestion mixture was diluted again with the same dilution buffer to reach 0.5 M GdmCl, followed by tryptic digestion (Roche, enzyme to protein ratio 1:50, MS-grade) and incubation at 37°C overnight in a thermal shaker at 700 rpm. Peptide desalting was performed according to the manufacturer’s instructions (Pierce C18 Tips, Thermo Scientific, Waltham, MA). Desalted peptides were reconstituted in 0.1% formic acid in water and further separated into four fractions by strong cation exchange chromatography (SCX, 3M Purification, Meriden, CT). Eluates were first dried in a SpeedVac, then dissolved in 5% acetonitrile and 2% formic acid in water, briefly vortexed, and sonicated in a water bath for 30 seconds prior injection to nano-LC-MS/MS.

#### Run parameters

LC-MS/MS was carried out by nanoflow reverse-phase liquid chromatography (Dionex Ultimate 3000, Thermo Scientific) coupled online to a Q-Exactive HF Orbitrap mass spectrometer (Thermo Scientific), as reported previously (Ni et al., 2019). Briefly, the LC separation was performed using a PicoFrit analytical column (75 μm ID × 50 cm long, 15 µm Tip ID; New Objectives, Woburn, MA) in-house packed with 3-µm C18 resin (Reprosil-AQ Pur, Dr. Maisch, Ammerbuch, Germany).

#### Peptide analysis

Raw MS data were processed with MaxQuant software (v1.6.10.43) and searched against the mouse proteome database UniProtKB with 55,153 entries, released in August 2019. The MaxQuant processed output files can be found in Table S8, showing peptide and protein identification, accession numbers, % sequence coverage of the protein, q-values, and label-free quantification (LFQ) intensities. The mass spectrometry data have been deposited to the ProteomeXchange Consortium (http://proteomecentral.proteomexchange.org) via the PRIDE partner repository (Martens et al., 2005) with the dataset identifier PXD026641. Complete proteome profiles are available in Table S3.

#### Differential expression analysis

The DEP package (version 1.6.1) was used. First, duplicate proteins, contaminants, and proteins that were not found in at least 2 out of the total number of samples (n=6) were filtered. A total number of 4,260 proteins were identified. The LFQ values were normalized using the background correction variance stabilizing transformation (VSN). Missing values were imputed using a left-censored imputation method as the proteins with missing values were biased to low expression values. Differential expression analysis was run with test_diff function from DEP package which uses limma (Ritchie et al., 2015). Differentially expressed proteins (DEP) were defined applying multiple-testing adjusted P (Benjamini-Hochberg) < 0.05 significant threshold and fold-change with an absolute value of > 1.5. Complete lists of DEP are available in Table S4.

#### Scatter plots

show mean LFQ values in hypoxia vs normoxia and were generated using ggplot2. GO-BP analysis was performed using DAVID web tools (Dennis et al., 2003).

### Simultaneous determination of cytidine modifications by targeted LC-MS/MS

#### Sample preparation

Genomic DNA was extracted using the PureLink Genomic DNA Mini Kit following the manufacturer ‘s instructions. 300 ng genomic DNA were used for profiling the following cytidine modifications in E14 mouse ES cells samples: 2′-deoxycytidine (dC), 5-methyl-2′-deoxycytidine (5-mdC), 5-hydroxymethyl-2′-deoxycytidine (5-hmdC), 5-formyl-2′-deoxycytidine (5-fodC), and 5-carboxyl-2′-deoxycytidine (5-cadC), as well as the other three bases 2′-deoxyguanosine (dG), 2′-deoxyadenosine (dA), thymidine (T). A multiple reaction monitoring (MRM) method with three transitions was established by using pure compounds. Furthermore, a dilution series of these standards was used for absolute quantification. 300 ng DNA was dissolved in a total of 21.5 µL water in Protein LoBind Tubes, 2.5 µL DNA Degradase buffer, and 1 µL DNA Degradase Plus enzyme (Zymo Research, Freiburg, Germany) were added. The DNA digestion efficiency was monitored by using 100 ng 5-methylcytosine and 5-hydroxymethylcytosine DNA standard sets (Zymo Research) in parallel under identical conditions. The digest was carried out at 37°C on a rocking platform (600 rpm) for 2 hours. 1 µL of 100 µM chloramphenicol was added as an internal standard as well as 1 µL of 5% formic acid to inactivate the digest.

#### Run parameters

180 ng of the DNA digests were used for LC-MS/MS analysis. Cytidine separation was performed on an LC instrument (1290 series UHPLC; Agilent, Santa Clara, CA), online coupled to a triple quadrupole hybrid ion trap mass spectrometer QTrap 6500 (Sciex, Foster City, CA). Cytidines were eluted from a Reprosil-PUR C18-AQ (1.9 μm, 120 Å, 150 × 2 mm ID; Dr. Maisch; Ammerbuch, Germany) column at a controlled temperature of 30 °C, using a gradient from 2 to 98% solvent B in solvent A over 10 min at 250 µL per minute flow rate. Solvent A was 10 mM ammonium acetate, pH 3.5 (adjusted with acetic acid) and solvent B was 0.1% formic acid in acetonitrile. Transition settings are provided in Table S9, consisting of three transitions for each base. Transitions were monitored in a 240-second window of the expected elution time and acquired at unit resolution (peak width at 50% was 0.7 ± 0.1 Da tolerance) in quadrupole Q1 and Q3. Data acquisition was performed with an ion spray voltage of 5.5 kV in positive mode of the ESI source, N2 as the collision gas was set to high, curtain gas was set to 30 psi, ion source gas 1 and 2 were set to 50 and 70 psi, respectively, and an interface heater temperature of 350 °C was used.

#### Analysis

Relative quantification of the peaks was performed using MultiQuant software v.2.1.1 (Sciex, Foster City, CA). The integration settings were a peak-splitting factor of 5 and a Gaussian smoothing width of 2 All peaks were reviewed manually. Only the average peak area of the first transition was used for calculations. Normalization was done according to IS. All original LC-MS generated QTrap wiff files, as well as MultiQuant v.2.1.1 (Sciex, Foster City, CA). Results were plotted using GraphPad Prism.

### ChIP-sequencing

Chromatin extracts were prepared as described in (Brookes et al., 2012). Briefly, cells were fixed with 1% formaldehyde (Thermo, 28906) in PBS at RT for 10 minutes. Then, 0.125M glycine (Sigma, 50046) was added to quench formaldehyde at RT for 5 minutes. Cells were washed twice with ice-cold PBS. To lyse, fixed cells were treated with swelling buffer (25mM HEPES pH 7.9, 1.5mM MgCl_2_, 10mM KCl (Invitrogen, AM9640G), 0.1% Igepal 630, 1x protease inhibitor cocktail, 1mM PMSF, 2mM NaVO_3_, 5mM NaF) at 4°C for 10 minutes. Cells were scraped on ice and passed through 18G needles before centrifugation at 3000g, 4°C for 5 minutes. Cell pellet, corresponding to nuclei was carefully resuspended in sonication buffer (50mM HEPES pH 7.9, 140mM NaCl, 1mM EDTA, 1% Triton X-100, 0.1% Sodium-deoxycholade (Thermo, 89904), 0.1% SDS (Invitrogen, AM9822), 1mM PMSF, 2mM NaVO_3_, 5mM NaF) and incubated on ice for 10 minutes before sonication. Chromatin was sheared to an average size of 200-300bp with E220 Evolution Covaris sonicator for 6 cycles, 1 minute each. Shearing efficiency was checked by agarose gel.

10µg chromatin was incubated with 5µL of HIF1α antibody (concentration not provided) (CST, 36169, lot:2), 20 µL of Protein A/G dynabeads (Thermo, 10002D/10004D) in sonication buffer overnight. Beads were washed twice with sonication buffer, followed by a wash with high salt buffer (sonication buffer with 500mM NaCl instead) and TE buffer (10mM Tris-HCl pH 8.5, (Teknova, T5085), 1mM EDTA, 1% SDS). Then, the sample was resuspended in elution buffer (50mM Tris-HCl pH 7.5 (Sigma, T2319), 1mM EDTA, 1% SDS). Samples were treated with RNase A (Thermo, EN0531) at 37°C for 30 minutes, followed by Proteinase K (NEB, P8107S) at 65°C overnight. Genomic DNA was purified using MinElute PCR purification kit (Qiagen, 28004) and quantified by Qubit 3.0. Two biological replicates were collected per condition.

#### Library preparation and sequencing

Libraries were prepared using the KAPA Hyper Prep Kit (Kapa Biosystems, KR0961) (input amount 10ng DNA, adapter concentration 1.5 µM, and size selection of 200-700-bp after PCR with cycle number = 15). Samples were sequenced using NovaSeq 6000, in 100-bp, paired-end format.

#### Mapping and analysis

Raw reads of treatment and input samples were subjected to adapter and quality trimming with cutadapt (Martin, 2011) (version 2.4; parameters: --nextseq-trim 20 --overlap 5 --minimum-length 25 --adapter AGATCGGAAGAGC -A AGATCGGAAGAGC).

Reads were aligned to the mouse genome (mm10) using BWA with the ‘mem’ command (version 0.7.17, default parameters). A sorted BAM file was obtained and indexed using samtools with the ‘sort’ and ‘index’ commands (version 1.10). Duplicate reads were identified and removed using GATK (version 4.1.4.1) with the ‘MarkDuplicates’ command and default parameters. Peaks were called with reads aligning to the mouse genome only using MACS2 ‘callpeak’ (version 2.1.2; parameters -- bdg --SPMR) using the input samples as control samples. For downstream analyses, after validation of reproducibility, replicates were pooled using samtools ‘merge’. Genome-wide coverage tracks (signal files) for merged replicates normalized by library size were computed using samtools bamCoverage (version 3.4.3) (parameters: --normalizeUsing CPM --extendReads) When required, bedgraph files were also generated using bigWigToBedGraph from Kent utils tools (Kent et al., 2010). For identification of consistent HIF1α peaks, only those that were identified in both replicates were used for downstream analyses. Peaks were annotated using ChIPseeker package (version 1.20.0) using default parameters (TSS region ±3-Kb) and subdivided into peaks at promoter or distal regions. Complete list of HIF1α peaks is available in Table S5.

Density plots were generated using computeMatrix (reference-point -- referencePoint center) and plotHeatmap functions from Deeptools. Enrichment of different features was at ±5-Kb of centered HIF1α peaks at promoters or distal regions. H3K27ac, p300, H3K4me3 and H3K4me1 ChIP-seq data generated on E14 ES cells were retrieved from (Cruz-Molina et al., 2017). Genome view of selected loci were generated using SparK (version 2.6.2). β-catenin ChIP-seq data generated on V6.5 ES cells was retrieved from (Zhang et al., 2013) and analyzed following our ChIP-seq workflow.

Overlap of Hif1α-, β-catenin- and ES enhancer peaks was generated using intersect function (-wa -wb) from Bedtools (version 2.29.2). Venn diagram was generated manually.

#### Density plots

were generated using compute Matrix (reference-point -- referencePoint) for TSS and (scale-regions) for gene body resolution and plotHeatmap functions from Deeptools. Enrichment of different features was defined at 1) ±2.5-Kb of DE genes at promoters identified by RNA-seq.

### WGBS

#### Sample preparation

Genomic DNA was extracted from 1 million cells using the PureLink Genomic DNA Mini Kit (Thermo, K182002) following manufacturer’ s instructions. gDNA was sheared in Covaris micro TUBE AFA Fiber Pre-Slit Snap-Cap tubes (SKU, 520045) and purified with the Zymo DNA Clean & Concentrator-5 kit (D4013) according to manufacturer’s instructions. The purified DNA was then bisulfite converted using the EZ DNA Methylation-Gold Kit (Zymo, D5005).

#### Library preparation and sequencing

WGBS libraries were processed using the Accel-NGS Methyl-seq DNA library kit (Swift Biosciences, DL-ILMMS-12) following manufacturer’s recommendations. Libraries were prepared and cleaned using Agencourt AMPure XP beads (Beckman Coulter, A63881). The absence of adapters from the final libraries was verified using the Agilent TapeStation. The WGBS libraries were sequenced on the NovaSeq6000, in 150-bp, paired-end format.

#### Mapping and analysis

Raw reads were subjected to adapter and quality trimming with cutadapt (version 2.4; parameters: --quality-cutoff 20 --overlap 5 --minimum-length 25; Illumina TruSeq adapter clipped from both reads) followed by trimming of 10 nucleotides from the 5’ and 3’ end of both reads. The trimmed reads were aligned to the mouse genome (mm10) using BSMAP (version 2.90; parameters: -v 0.1 -s 16 -q 20 -w 100 -S 1 -u-R). Duplicates were removed using the ‘MarkDuplicates’ command from GATK (version 4.1.4.1; --validation_stringency=lenient --remove_duplicates=true). Methylation rates were called using mcall from the MOABS package (version 1.3.2; default parameters). Only CpGs covered by at least 10 and at most 150 reads located on autosomes or chromosome X were considered for all downstream analysis. Replicates were merged by calculating the average for each CpG covered by at least one of the two samples.

##### Genomic feature annotations

Genomic tiles were defined by segmenting the genome into regions of one kb size. The annotation of CGIs was downloaded from UCSC for the mouse reference genome mm10. CGI shores were defined as the two kb regions flanking each CGI on either side. CGI shelves were defined as the two kb flanking the outer sides of CGI shores. Annotation of repeat elements was downloaded from RepeatMasker. The gene annotation was downloaded from GENCODE (VM19) and promoters were defined as the regions 1.5 kb upstream and 500 bp downstream of the TSS.

##### Feature-wise DNA methylation

Average methylation levels per feature were calculated using the arithmetic mean based on the CpGs commonly covered by all samples (merged replicates). Only features with at least three covered CpGs were considered. For methylation differences per feature, the arithmetic mean of one sample was subtracted by the arithmetic mean of another sample.

Heatmaps of average promoter methylation were prepared using the ‘pheatmap’ R package. Promoters were classified as overlapping with a CGI if at least 20% of the promoter or CGI were overlapping with each other.

##### DNA methylation levels at HIF1α peaks

HIF1α peaks at both promoter and distal regions were merged. 1-Kb tiles signal files were used to quantify the mean methylation levels at HIF1α peaks using map function (-c 4 - mean) from Deeptools. Mean methylation levels were used to generate violin plot using ggplot2.

### Gastruloid formation

Male F1G4 mouse ES cells (George et al., 2007) were cultured on 6cm plates (Corning 430166) gelatinized with 0.1% gelatin (Sigma G1393) and coated with mitotically inactive primary mouse embryo fibroblasts (3-4×10_4_ cells/cm^2^) with standard ES medium containing 15% FCS and 1000 U/ml leukemia inhibitory factor (LIF, Chemicon ESG1107) at 37°C and 5% CO2. ES cells were split every second day with a 1:10 dilution. For splitting, media was aspirated and cells were washed once with PBS and trypsinized (Trypsin-EDTA (0.05%) (Gibco 25300054)) for 5-10 minutes at 37°C. Trypsin was neutralized by 3 ml ES media and cells centrifuged for 5 minutes at 1000 rpm, after which the pellet was resuspended in ES media. ES cells were cultured in normoxia or hypoxia for 7 days. Gastruloids were then generated as described previously (Veenvliet et al., 2020) with some minor modifications. Briefly, ES cells were first depleted of feeders, then washed once in 5ml pre-heated (37°C) PBS containing MgCl2 and CaCl2 (Sigma D8662) and once in 5ml NDiff227 medium (Takara Y40002) pre-conditioned in normoxia or hypoxia. ES cells were then pelleted by centrifugation for 5 minutes at 1000rpm and resuspended in 250µl of NDiff227. 10µl of the cell suspension was mixed with 10µl of Trypanblue (Bio-Rad 1450021) for automated cell counting with Luna Automated Cell Counter. 400 live cells were plated in a volume of 35µl NDiff227 into each well of a 96-well round bottom, low attachment plate (Costar 7007 ultra-low attachment 96 well plate (7007)). Cells were then allowed to aggregate for 48 h under normoxic or hypoxic conditions. After 48h, the aggregates were treated with 3µM CHIR99021 (CHIR, Merck Millipore) in 150µl NDiff227 for 24h to induce robust gastruloid formation. For “-Chi” aggregates, 150µl NDiff227 without CHIR was added. Between 72 and 120h, medium was refreshed every 24h by removing 150µl of the old media and adding the same volume of new, pre-conditioned NDiff227.

### Whole-mount immunofluorescence (WIFC) of gastruloids

Gastruloids were picked using a p200 pipette with the tip cut-off at the 50µl mark. Gastruloids were washed twice with PBS + MgCl_2_ and CaCl_2_ + 0.5% BSA (Sigma A8412), once with PBS, and then fixed in 4% PFA for 75 minutes in Ibidi 8-well glass-bottom plates (Ibidi 80827) at 4°C on a rocking platform. Subsequently, gastruloids were washed twice in PBS for 5 min, permeabilized by incubating for 3 × 10 minutes in 0.5% Triton-X/PBS (PBST), and blocked in 10% fetal calf serum/PBST (blocking solution) overnight at 4°C. Primary antibody incubation was performed in blocking solution for 48-72h at 4°C, after which gastruloids were washed three times with blocking solution and three times with PBST. The following primary antibodies were used: rabbit anti-T (Cell Signaling, D2Z3J; 1:500), goat anti-SOX2 (R&D Systems, AF2018; 1:500), goat anti-FOXA2 (Santa Cruz, sc-6554; 1:500). After the last washing step, gastruloids were incubated in blocking solution o/n at 4°C. The next day, secondary antibodies diluted in blocking solution were added, and gastruloids were incubated for 24h at 4°C. The following secondary antibodies were used, all at a dilution of 1:500: donkey anti-rabbit Alexa Fluor 647 (Thermo, A31573), donkey anti-goat Alexa Fluor 546 (Thermo, A11056). Afterwards, gastruloids were washed three times with blocking solution and three times with PBST. The last PBST washing step after secondary antibody incubation included DAPI (0.02%, Roche Diagnostics 10236276001). DAPI was incubated for 5 hours or overnight and washed off once with PBS.

### Clearing and imaging of gastruloids

Prior to imaging, gastruloids were embedded in agarose and cleared with RIMS (Refractive Index Matching Solution). To this end, 1.5% low melting point (LMP), analytical grade (Promega, V2111) agarose was prepared in PBS, incubated at 80°C for 15 minutes, and cooled down to 37°C for 3 minutes in a thermomixer. Samples were washed twice with PBS for 10 minutes, post-fixed in 4% PFA for 20 minutes, and washed three times with 0.1M phosphate buffer (PB, 0.025M Na_2_HPO_4_, and 0.075M Na_2_HPO_4_, pH 7.4). The pipette was set to 20µl and the gastruloids were stabilized on the IBIDI plate with a drop of LMP agarose for 5 minutes until the agarose was dry. The clearing was performed by incubation in 200µl RIMS (133% w/v Histodenz (Sigma, D2158) in 0.02M PB) on a rocking platform at 4°C for at least one to several days. Cleared gastruloids were imaged with the Zeiss LSM 880 Airyscan in confocal, Airyscan or Fastairy mode, using a Plan-Apochromat 20x/ NA=0.8 Objective, lateral pixel size of 0.1.2-1.2 µm and a typical z-spacing ranging from 1.9-3.3 μm and appropriate laser/filters for DAPI, Alexa Fluor 546, and Alexa Fluor 633 or combinations thereof. Bright-field images of non-cleared gastruloids were acquired with the Zeiss Celldiscoverer 7, with a Plan-Apochromat 5x/ NA=0.3 Objective and a 1x postmagnification, lateral pixel size of 0.9 µm and a typical z-spacing of 10 μm, running under ZEN blue v3.1.

### Post-acquisition image processing and analysis

Images captured in airy mode were processed under ZEN black 2.3 on a dedicated workstation. Confocal and wide-field image sets were downstream analyzed and further processed using a ZEN blue workstation (version 3.2). Maximum intensity projections (MIP) for gastruloids morphometrics analysis were achieved using customized macros in the open application development module in ZEN blue v3.2. Morphometric analysis was performed by either variance-based thresholding (bright-field images) or within confocal MIPs primary using Otsu intensity thresholds after faint gaussian smoothing, close by objects were segmented downstream by standard shedding.

Single cells image analysis of confocal datasets were also performed with customized analysis pipelines written in the image analysis module in ZEN blue. Briefly individual cells were identified by nuclear counterstainings after gaussian smoothing and background subtraction, adjusted to actual resolution of individual datasets, close by objects were segmented by water shedding. All objects/ nuclei were filtered after identification by area of 100-1000 µm^2^ and circularity of 0.6-1 (sqrt (4 × area / π × FeretMax^2^)). Within the resulting regions of interest (cells) fluorescence signals of the counterstaining as well as T/FOXA2/ SOX2 were quantified. In total, > 10 million single objects were analyzed, plotted and further quantified using customized R-scripts.

### Quantification of T expression across the anterior-posterior axis

T expression along the anterior-posterior axis was performed in ZEN 3.3 lite. In detail, Maximum intensity projections (MIP) previously developed by ZEN blue v3.2. were loaded into ZEN 3.3 lite. Brightness/contrast were automatically adjusted and a line (stroke thickness: 1) was manually drawn from the anterior to the posterior end of the structure along the midline, and the fluorescent intensity was measured using the “Profile” function on the software. The distance was measured in nm.

To obtained binned intensities, first relative positions were calculated by dividing the absolute position by the total length of the structure, resulting in standardized relative positions on a 0-1 scale. Average intensities were then calculated for 0.01 sized bins, resulting in average intensity for 100 bins over the anterior-posterior axis. Plotting and k-means clustering was performed using PlotTwist (Goedhart, 2020).

**Figure S1.**
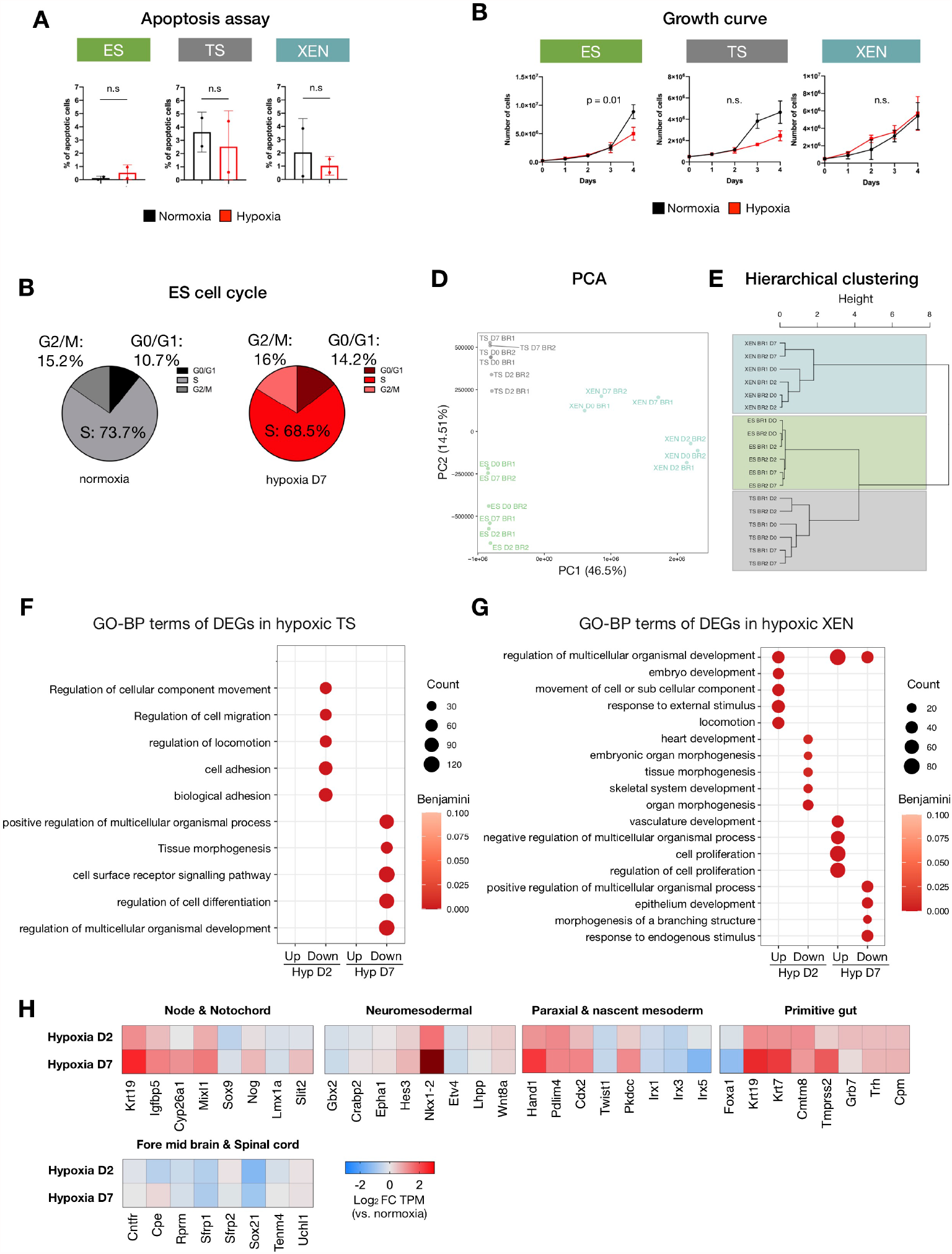
Further characterization of the hypoxic response in ES, TS, and XEN cells. A) Apoptosis levels of normoxic or hypoxic (d7) ES, TS, and XEN cells. Apoptotic cells were defined as Casp3/7^+^/Sytox^-^. See methods for more details. Two biological replicates were performed. The statistical test performed is a two-tailed paired Student’s t-test. B) Growth curves of ES, TS, and XEN cells grow in normoxia or hypoxia for 4 days. The same number of cells were seeded in both conditions and cells were counted daily. Two biological replicates were performed. The statistical test performed is a two-tailed paired Student’s t-test. C) Analysis of cell cycle phase distribution of normoxic or hypoxic (d7) ES cells. EdU and FxCycle Violet (DNA stain) were used. D, E) Principal component analysis (PCA) (d) and hierarchical clustering (e) based on global transcriptomes of ES, TS, and XEN cells in hypoxia day 2, hypoxia day 7, and normoxia. F, G) GO-BP terms associated with DE genes in TS (f) and XEN (g) cells exposed to acute (d2) or prolonged (d7) hypoxia. Representative significant terms are shown (Benjamini-Hochberg adjusted p-value < 0.1). No significant terms were retrieved for upregulated genes in TS cells on day 2 of hypoxia. See Table S2 for full list of GO terms. H) Heatmaps showing expression levels of the indicated genes in hypoxic relative to normoxic ES cells. Germ layer markers were retrieved by analyzing RNAseq data of gastrulating mouse embryos published in (Grosswendt et al., 2020) as explained in Methods (see ‘Selection of germ layers and lineage markers’ section). TPM, transcripts per million. FC, fold change.

**Figure S2.**
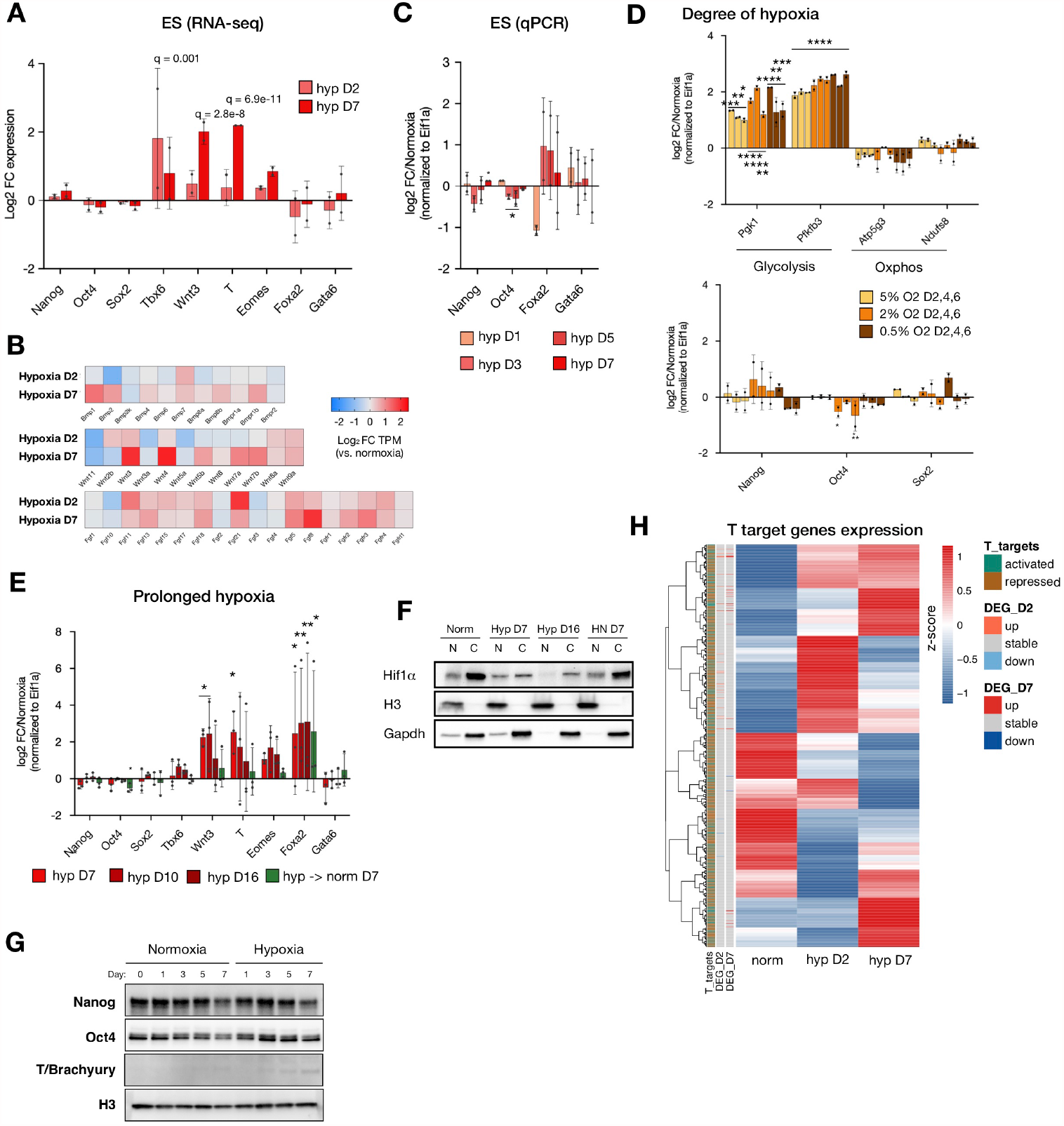
Further characterization of pluripotency and differentiation-associated gene activity in normoxic and hypoxic ES cells. A) Relative expression levels of shown genes measured by RNA-seq. Data represent log_2_FC over normoxic ES cells of TMP expression values and standard deviation. B) Heatmaps showing expression levels of Bmp, Wnt, and Fgf, genes in hypoxic relative to normoxic ES cells. C) Relative expression levels of shown genes in hypoxic ES cells measured by RT-qPCR. Data represent log_2_FC over normoxia normalized to Eif1α and standard deviation for two biological replicates. Other tested genes are available in Figure 2B. The statistical test performed is a two-tailed paired Student’s t-test. D) Relative expression levels of shown genes in ES cells exposed to different levels of hypoxia measured by RT-qPCR. Data represent log_2_ FC over normoxia normalized to Eif1α and standard deviation for two biological replicates. Other tested genes are available in Figure 2C. E) Relative expression levels of shown genes in ES cells exposed to different durations of hypoxia (2%) measured by RT-qPCR. Data represent log_2_ FC over normoxia normalized to Eif1α and standard deviation for three biological replicates. F) Western blot showing expression levels and subcellular localization of Hif1α in ES cells in normoxia or exposed to different durations of hypoxia. Gapdh and H3 were used as loading and fractionation controls. N, nuclear fraction. C, cytoplasmic fraction. HN D7, ES cells cultured in hypoxia for 7 days and then returned to normoxia for 7 days. G) Western blot showing expression levels of the indicated genes in normoxic and hypoxic ES cells. H3 was used as a loading control. H) Heatmap showing expression levels of T target genes in normoxic and hypoxic ES cells. T target genes were identified as described in the Methods (see ‘Analysis of T target genes’ section) by analyzing data from (Lolas et al., 2014)). Bar annotations show, 1) T_targets; T -activated (in green) and -repressed (in brown) target genes in *in vitro* primitive streak differentiated cells, 2) DEG in hypoxia day 2 (D2) and day 7 (D7); up-(in red), down-(in blue) regulated and stable (in gray) genes on hypoxia day 2 and/or day 7. Statistical tests performed are two-way ANOVA unless otherwise specified. P value style: p>0.05 (ns), p<0.05 (*), p<0.01 (**), p<0.001 (***), p<0.0001 (****).

**Figure S3.**
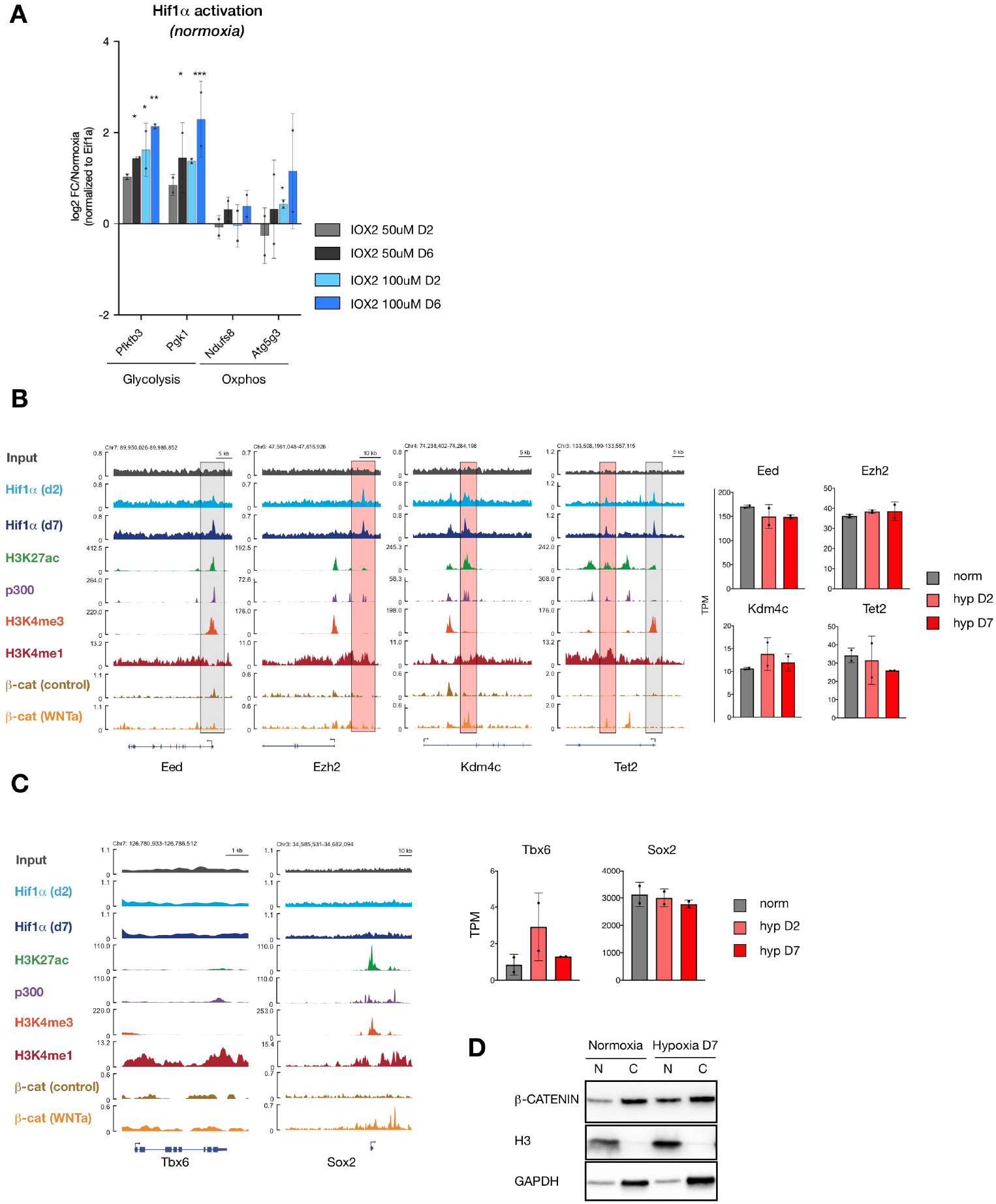
Further characterization of Hif1α activity at chromatin in hypoxic ES cells. A) Relative expression levels of shown genes in ES cells treated with the HIF1α activator IOX2 at indicated concentrations in normoxia. Data represent log_2_FC over DMSO-treated ES cells normalized to Eif1α and standard deviation for two biological replicates. Other tested genes are available in Figure 5A. Statistical test performed is a two-way ANOVA. B, C) Genome browser views of chromatin occupancy and histone modifications at and epigenetic regulators (b) and lineage-specific genes (c). Expression values of the shown genes in normoxic and hypoxic ES cells as measured by RNA-seq are plotted as bar plots (right panels). Gray and red highlights indicate promoter and ES active enhancers respectively. A) Western blot showing expression levels and subcellular localization of β-CATENIN in normoxic and hypoxic (d7) ES cells. Histone H3 is used as loading control. H3 and GAPDH were used as loading and fractionation controls. N, nuclear fraction. C, cytoplasmic fraction.

**Figure S4.**
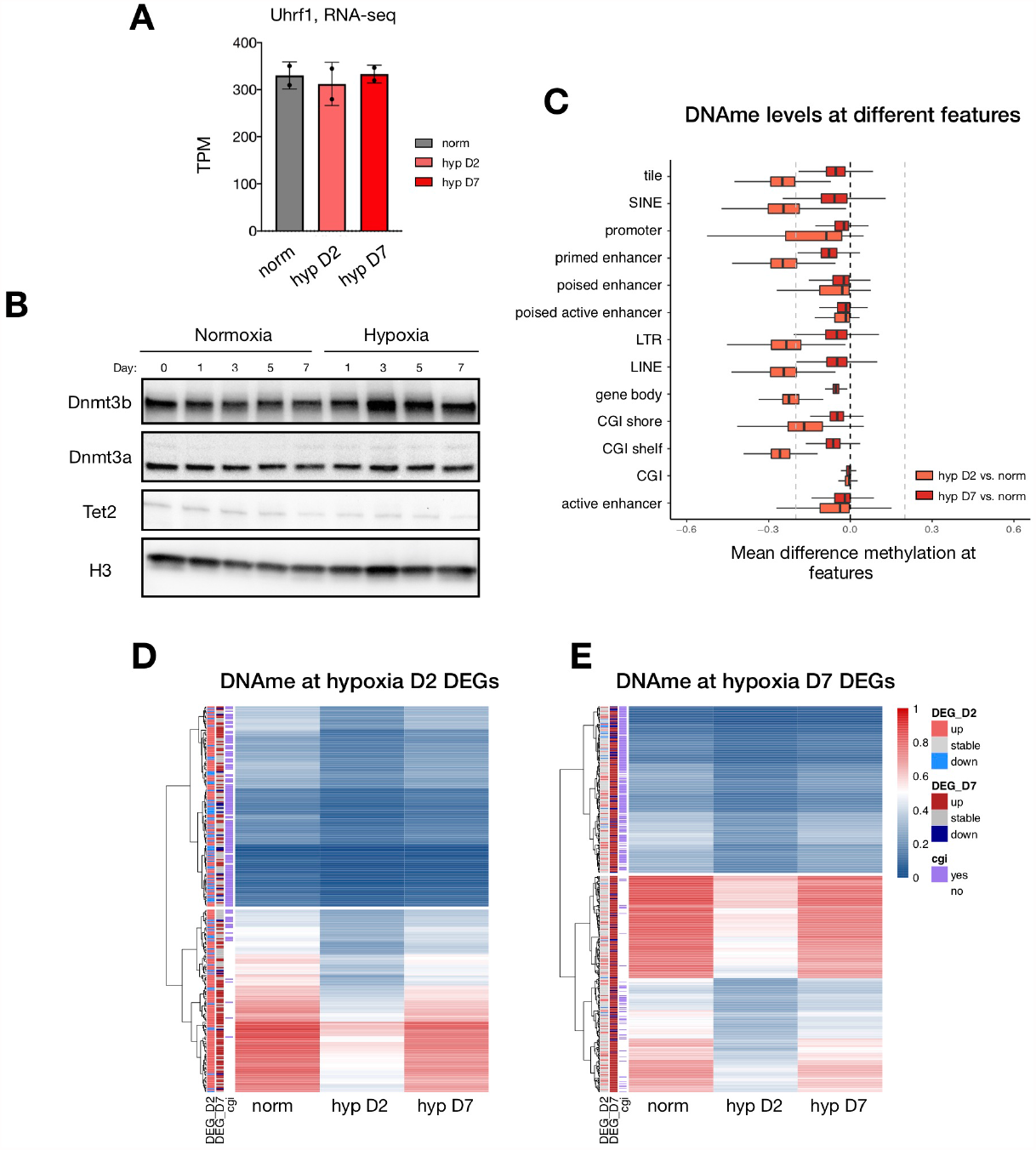
Characterization of DNA demethylation across the genome. A) Expression levels of Uhrf1 in normoxic and hypoxic ES cells as measured by RNA-seq. TPM, transcripts per million. B) Western blot showing expression levels of DNMT3b, DNMT3a, and TET2 in normoxic and hypoxic ES cells. H3 was used as loading control. C) Box plot panel showing levels of DNA methylation reduction at different genomic features. Data represents mean difference methylation compared to normoxia of two biological replicates. Line indicates median and whiskers indicate variability outside the upper and lower quartiles. D, E) Heatmaps showing mean DNA methylation levels of DE gene promoters identified in day 2 (c) or day 7 of hypoxia (d). Bar annotations (reported as DEG_D2 and DEG_D7) show the up-(in red), down-(in blue) regulated and stable (in gray) genes in hypoxia day 2 or day 7.

**Figure S5.**
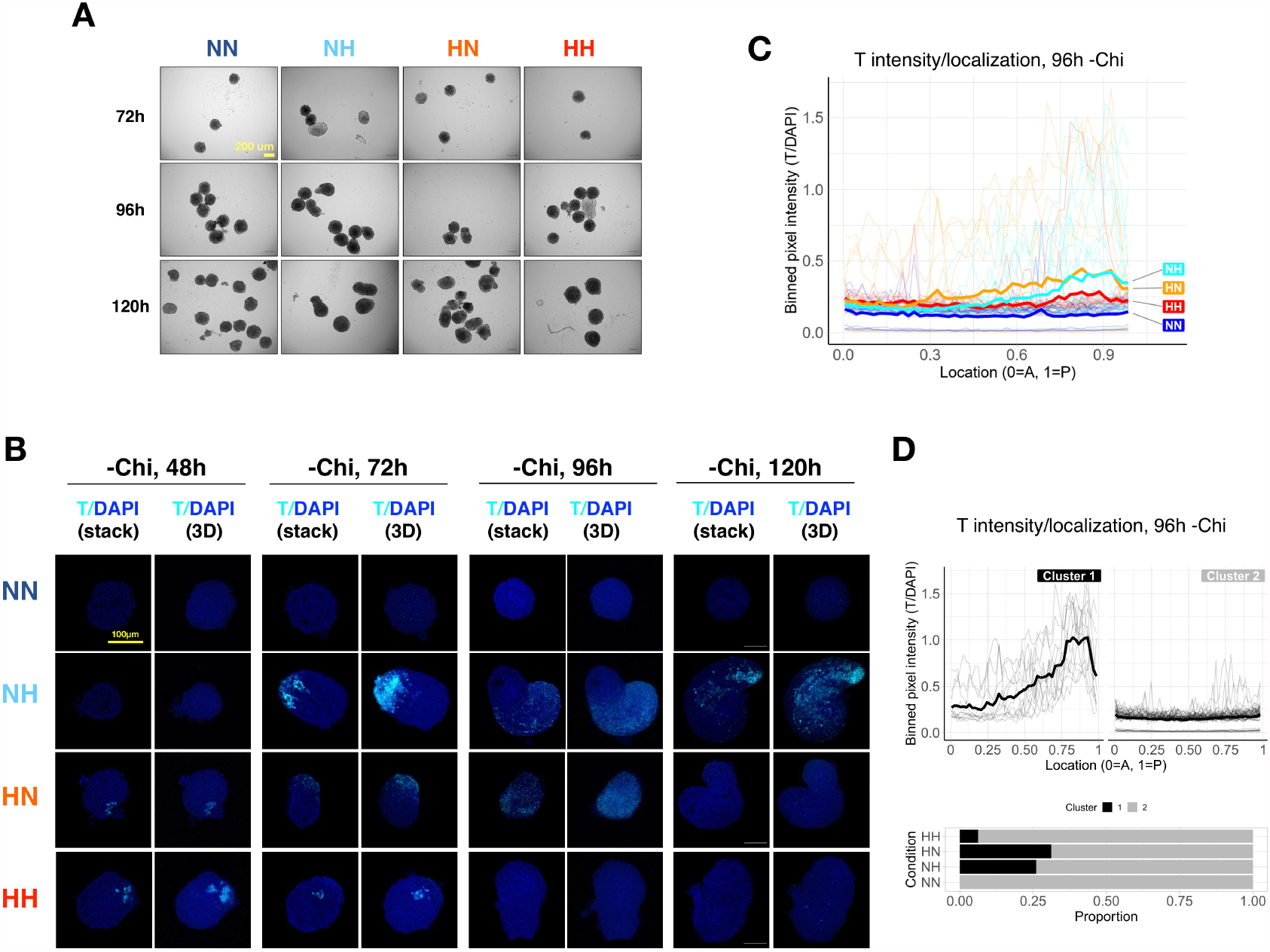
Hypoxia can induce spontaneous elongation of gastruloids in the absence of exogenous WNT activation. A) Bright-field images of representative gastruloids at the indicated time points. B) Confocal fluorescent microscopy images of representative -Chi gastruloids at indicated time points of culture. 3D, three-dimensional projection. Images were taken using the same settings for each time point, which may differ between time points. C) Localization of T signal along the anterior-posterior (A-P) axis of gastruloids in each condition at 96h of culture. T signal normalized to DAPI and binned at 1% length increments along each structure for plotting. See Methods for details. Thick lines show mean values and thin lines show data from individual structures. D) K-means clustering of the NH and HN structures presented in C) with n=2 clusters.

**Figure S6.**
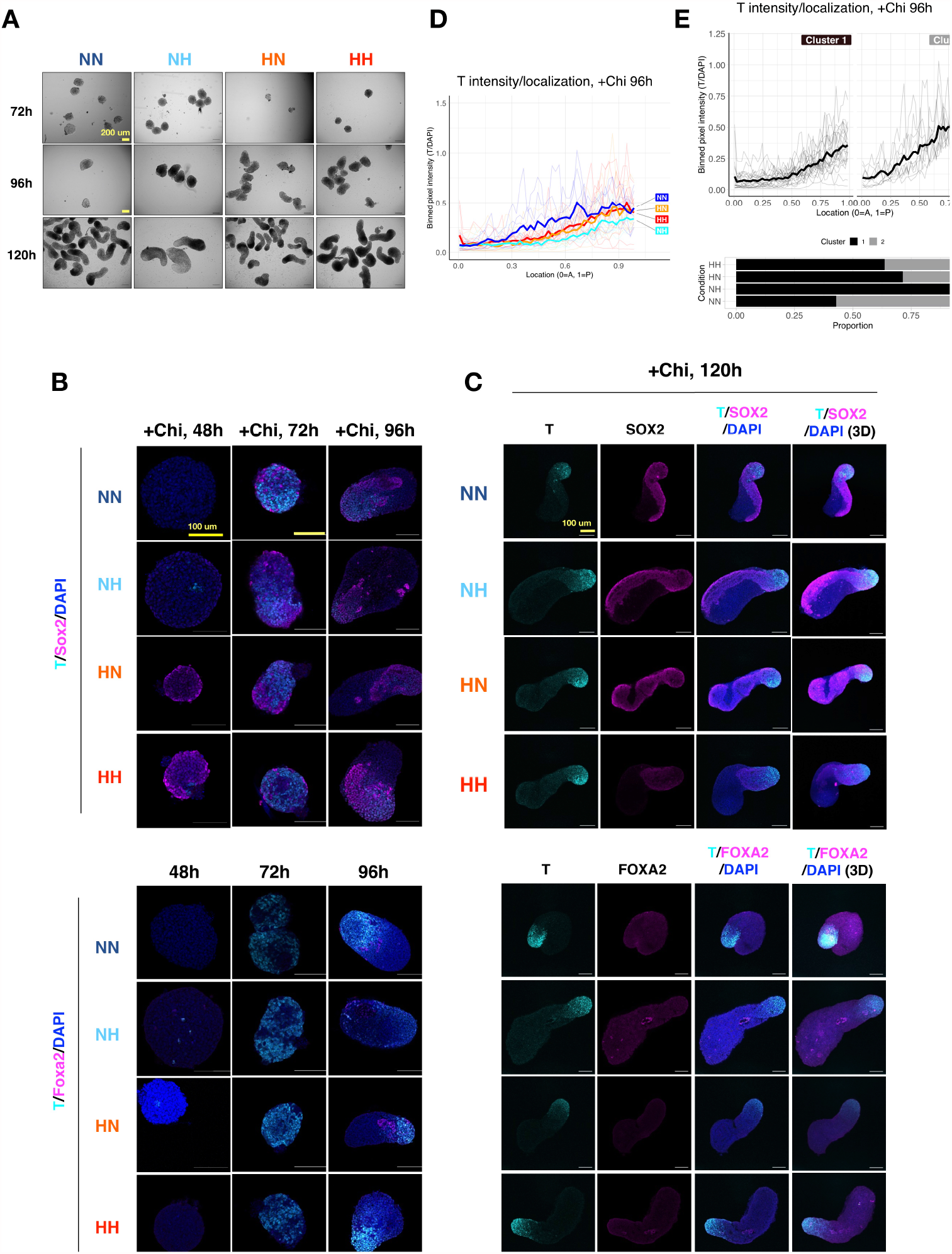
Further characterization of conventional gastruloids exposed to hypoxia. A) Bright-field images of representative gastruloids at the indicated time points. B) Fluorescent microscopy images of representative gastruloids at early time points. Images show a single z-stack. For the same time point, images were taken using the same settings across conditions, but settings differ between time points. C) Confocal fluorescent microscopy images of representative gastruloids at 120h of culture. Images show a single z-stack, except those labeled 3D, which show 3D maximum intensity projections of the structure. D) Localization of T signal along the anterior-posterior (A-P) axis of gastruloids in each condition. T signal was binned at 1% length increments along each structure for plotting and normalized to DAPI signal. Thick lines show the mean and thin lines show data from individual structures. E) K-means clustering of all 96h +Chi structures plotted in C) with n=2 clusters.

